# Glioblastoma cells imitate neuronal excitability in humans

**DOI:** 10.1101/2024.01.08.574637

**Authors:** Tong Tong, Josephine D. Hendriksen, Kirstine J. Elbæk, Attila Ozsvar, Jens T. Eschen, Francisco G. Rodríguez-González, Ilayda D. Pusat, Emilie Littau Christensen, Mads Rahbæk, Kathrine Pii Frederiksen, Søren O. S. Cortnum, Ann K. Sindby, Kaare Meier, Nikola Mikic, Jens C. H. Sørensen, Jonathan T. Ting, Marco Capogna, Bjarne W. Kristensen, Joachim Weischenfeldt, Wen-Hsien Hou, Anders R. Korshøj

**Author notes:** Deceased (Marco Capogna). **Corresponding authors:** Anders Rosendal Korshøj, MD, PhD, Aarhus Hospital, Department of Clinical Medicine, Neurosurgery, Palle Juul-Jensens Boulevard 165J, 8200, Aarhus N, Denmark.; Wen-Hsien Hou, PhD, Aarhus University, Department of Biomedicine, Høegh-Guldbergs Gade 10, 8000 Aarhus C, Denmark.

## Abstract

**Background:** Glioblastomas are renowned for their pronounced intratumoral heterogeneity, characterized by a diverse array of plastic cell types. However, the physiological and transcriptomic features of the cells residing in the invasive leading edge (LE), including both neurons and glioblastoma cells (GBCs), remain unclear, challenging our comprehension of the glioblastoma pathophysiology.

**Methods:** To elucidate molecular and morphophysiological features of LE cells, we established an experimental workflow enabling the investigation of GBCs and neurons within cancer-infiltrated organotypic tissue specimens from the same patients. With this approach, we characterized the electrophysiological properties of cells in the neocortical tumor LE (LE cells). We further performed single-cell Patch-seq experiments, enabling transcriptomic analysis of electrophysiologically recorded LE cells.

**Results:** Upon depolarization, 58% of LE cells exhibited aberrant action potentials (aAPs). Electrophysiological assessment showed that a subset of GBCs generated aAPs, with no significant differences in aAP properties compared to LE neurons. Transcriptomic analysis of 144 LE cells revealed four transcriptomic clusters, including two GBC populations and two neuronal populations. LE GBCs exhibited diverse cellular states, including mesenchymal-like, astrocyte-like, neural progenitor-like, and oligodendrocyte-precursor cell-like phenotypes. Notably, LE GBCs exhibiting aAPs displayed reduced mitotic pathway activity and developmental regulatory ion channel Ca_V_1.2. Cell-cell interaction analysis illustrates a higher signaling interaction between aAP LE cells compared to no-aAP LE cells.

**Conclusion:** In summary, we find comparable electrical properties between neurons and a subset of GBCs in the leading edge, suggesting an active electrophysiological role of GBCs in the tumor’s pathophysiology.

**Key Points:**

- Human organotypic slice cultures enable long-term functional investigation of glioblastoma cells.
- GBCs and neurons in the LE exhibit similar aAPs.
- LE GBCs display heterogeneous cellular states, with reduced proliferation signaling in aAP GBCs.

**Importance of study:** This study sheds light on the diverse pathophysiological and molecular features of cells in the neocortical infiltration zone of glioblastoma. By utilizing *in vitro* organotypic human brain slice cultures, a reliable platform for longitudinal observation, we unveiled the dichotomy of the electrical properties of LE cells. More than half of LE cells display aAP, contradicting findings from cultured tumor cells and animal models. Patch-seq analysis confirmed that both GBCs and neurons in the LE generate aAPs with indistinguishable electrical properties. aAP GBCs showed higher cell-cell interactions. We find aAP GBCs to express reduced proliferation signaling compared to no-aAP GBCs, suggesting a non-dividing and potentially more plastic cell state. These findings point to the electrical-active GBCs as an important attribute of the LE, linking the electrophysiological properties with functional implications, and implicate an active role of electrophysiological changes of GBCs in tumor pathophysiology.

## Introduction

Glioblastoma is a highly aggressive and incurable primary brain cancer^1^. These tumors display significant resistance to comprehensive treatment, leaving patients facing a dismal prognosis^2^. Glioblastomas are notably diverse in their cellular makeup^3–5^. In the tumor core, astrocytic (AC)-like and mesenchymal (MES)-like cell states form a resilient, self-repairing network, interconnected by multiple tumor microtubes and Connexin 43-dependent gap junctions^6–8^. In contrast, unconnected neural progenitor cell (NPC)-like and oligodendrocyte precursor cell (OPC)-like states are prevalent at the tumor’s infiltrative zone, moving outward from the core to colonize the surrounding brain^9–11^.

Despite these intriguing molecular features, the electrophysiological properties of GBCs at the LE remain largely elusive. Specifically, it is unclear if LE GBCs have excitable membranes, potentially enabling active integration into the surrounding tumor network and adjacent brain tissue. Preliminary analyses based on animal models and glioblastoma cell lines have indicated a non-excitable nature of GBCs^12–14^, akin to astrocytes^15^. Conversely, one study on acute human glioblastoma-infiltrated brain slices reported action potentials (APs) in putative tumor cells^16^. However, it was unclear whether these cells represented GBCs or non-neoplastic cells. By a similar notion, a few studies have reported that some low-grade oligodendroglioma and astrocytoma cells generate aberrant APs [17, 18].

These discrepant observations highlight the need for elucidation of the intrinsic electrophysiological and genetic profiling of cells in the complex glioblastoma LE. One of the challenges is to tackle the difficulty of identifying cellular subtypes within the heterogeneous tumor microenvironment, as these can be elusive under acute conditions. In addition, the current understanding of the physiological attributes of GBCs derives mainly from animal models and cell lines, limiting generalizability to humans and highlighting the need for human-based model systems.

To overcome this, we utilized an approach based on stable human organotypic slice cultures to investigate cells in the glioblastoma neocortical LE. This approach enabled us to identify and characterize the electrophysiological, morphological, and transcriptomic properties of both GBCs and non-neoplastic cells at the tumor LE (LE cells), as well as neurons in the adjacent neocortex (non-LE neurons). Multimodal cellular signatures were obtained by patch-seq, an approach combining whole-cell patch-clamp recordings with single-cell transcriptomic profiling. This method enables direct correlation between electrophysiological phenotype and gene expression within the same cell, providing critical insight into the heterogeneous landscape and functional importance of GBCs and non-neoplastic cells in the LE.

## Materials and Methods

### Human glioblastoma tissue acquisition

Human tissue procedures were approved by the Central Denmark Region Committee for Health Research Ethics (Journal Number: 1-10-72-82-17) and adhered to the World Medical Association Declaration of Helsinki principles^19^. With patient consent, cortical tumor-infiltrated tissue was resected from 18 glioblastoma patients at Aarhus University Hospital, with diagnoses confirmed by WHO 2021 criteria (Supplementary Fig. 1)^1^. Preoperatively, patients received 5-Aminolevulinic acid hydrochloride (5-ALA) or sodium fluorescein to visualize the tumor-infiltrated areas for sampling and resection^20, 21^. The resected tissue was taken from cortical transition zones between contrast-enhancing and non-enhancing regions on T1-weighted MRI, retaining intact pia mater and showing 5-ALA or fluorescein fluorescence during microsurgical resection. Tissue specimens, ranging from 1-2 cm^3^, were resected and immediately immersed in ice-cold artificial cerebrospinal fluid (ACSF) containing (in mM): 75 Sucrose, 84 NaCl, 2.5 KCl, 1 NaH_2_PO_4_, 25 NaHCO_3_, 25 D-glucose, 0.5 CaCl_2_·2H_2_O and 4 MgCl_2_·6H_2_O, with a pH range of 7.3-7.4. The samples were transported to the laboratory in carbogenated ACSF (95% O_2_, 5% CO_2_) within 30 minutes.

### Acute and cultured human organotypic slice preparation

The human ex vivo brain slice culture method was adapted from established methods [22-24]. 350-μm-thick slices were cut in ice-cold carbogenated ACSF using a vibratome (Leica 1200S). After a 30-minute recovery in carbogenated ACSF at 34°C, slices were transferred to ACSF containing (in mM): 130 NaCl, 3.5 KCl, 1 NaH_2_PO_4_, 24 NaHCO_3_, 10 D-glucose, 1 CaCl_2_·2H_2_O, and 3 MgCl_2_·6H_2_O with a pH range of 7.3-7.4, at room temperature. The slices were then divided either for acute recordings or cultured on membrane inserts in 6-well plates for virus transduction. The slice culture media contained (in mM): 8.4 g/L MEM Eagle medium, 20% heat-inactivated horse serum, 30 HEPES, 13 D-glucose, 15 NaHCO_3_, 1 ascorbic acid, 2 MgSO_4_·7H_2_O, 1 CaCl_2_.4H_2_O, 0.5 GlutaMAX-I and 1 mg/L insulin, 25 U/mL Penicillin/Streptomycin, with a pH range of 7.2–7.3 adjusted by Tris-base, and osmolality of 295–305 mOsm/kg, and sterile-filtered. Medium was refreshed every 1-2 days until use.

### AAV-viral transduction

The adeno-associated virus (AAV) vectors were acquired from the University of Zurich Viral Vector Facility. AAV-hGFAP-EGFP (AAV5-hGFAP-hHEbl/E-EGFP-bGHp(A)) and AAV-hSyn1-EGFP (AAV1-hSyn-EGFP-WPRE-hGHp(A)) were used in this study. Viral titers ranged between 1–3 × 10^13^ units/ml. The viral vectors were applied to the slice surface using a fine micropipette at least one hour after the initial plating to allow the slice to equilibrate.

### Whole-cell patch-clamp recording

Tumor-infiltrated brain slices were placed in the recording chamber of a SliceScope microscope (Scientifica) and perfused with carbogenated ACSF at 32–35°C. A SciCam Pro camera (Scientifica) was used for visualization and image capture. Whole-cell recordings were performed using pulled borosilicate pipettes (5–7 MΩ; Harvard glass capillaries, GC120F-10) filled with an internal solution containing in (mM) 126 K-gluconate, 10 HEPES, 4 Mg-ATP, 0.3 Na_2_-GTP, 4 KCl, 10 phosphocreatine, 8 biocytin, and 50 μM Alexa-594 with a pH range of 7.2–7.3 and 295 mOsm/kg osmolality. Data were acquired with a Multiclamp 700B and Digidata 1550B (Molecular Devices, US) and were low-pass filtered at 2 kHz and digitized at 20 kHz. Cells were accepted if they had >1 GΩ seal resistance and access resistance <20 MΩ. Resting membrane potential (RMP) was measured after break-in, and APs were evoked by 3-ms depolarizing steps. aAPs were defined by a derivative threshold (>20 mV/ms)^25^ and quantified for initiation, amplitude, half-width, and 10–90% rise/decay slopes. To confirm ionic contributions, TEA (30 mM, Sigma-Aldrich) and TTX (1 µM, TOCRIS) were sequentially applied. Coordinates of recorded cells were saved for later histology.

### Nucleus collection following patch-clamp recordings

At the end of the recordings, nuclei were extracted by gentle negative pressure, aspirated into the pipette, and expelled into the lysis buffer with RNase inhibitor (Takara)^24^. Samples were snap-frozen on dry ice and stored at –80 °C for transcriptomic processing.

### cDNA amplification, library construction and sequencing

Following nucleus collection, reverse transcription and cDNA amplification were performed using the SMART-seq v2 Ultra Low Input RNA Kit (Takara). The cDNA quality and yield were assessed via Bioanalyzer (Agilent Technologies). Libraries were prepared using the Nextera XT DNA Library Preparation Kit (Illumina) and sequenced on Illumina NextSeq 500.

### RNA-seq gene expression quantification

Reads were aligned to GRCh38 using STAR v2.7.0c, with the parameters outFilterScoreMinOverLread=L0.3, outFilterMatchNminOverLread=L0.3, outFilterMatchNmin=0 and outFilterMismatchNmax=2 options. Gene counts were computed using the R Genomic Alignments summerizeOverlaps function with the “IntersectionNotEmpty” option.

### Transcriptomic data analysis

The count matrices were imported into R (version 4.1.0) and analyzed using Seurat (version 4.1.1)^26^. The cells underwent quality control and filtering, removing cells with more than 10% of transcripts mapping to mitochondrial genes or cells with less than 500 unique genes detected, resulting in a dataset of 144 cells. To distinguish between GBCs and non-neoplastic cells, CopyKat (version 1.1.0) and InferCNV (version 1.22.0, (inferCNV of the Trinity CTAT Project. https://github.com/broadinstitute/inferCNV were used to detect copy-number alterations (CNAs) in the cells. Consensus of cells showing CNA were annotated as GBCs, while the remaining cells underwent cell typing using label transfer with the glioblastoma Darmanis et al. dataset as reference^27^. Integration across patients was performed using Harmony^28^. For visualization, Uniform Manifold Approximation and Projection (UMAP) was calculated on the top 30 principal components. The GBCs were classified using the cellular states from Neftel et al.^4^ using AddModuleScore from Seurat, calculating the average expression level of each program subtracted by the average expression of a control gene set.

### Differential gene expression analysis

Differential gene expressions between LE aAP and no-AP (both neurons and GBCs) were assessed using the MAST test implemented in Seurat’s FindMarkers function. Cells were subset to the tumor population, and the patient of origin was included as a latent variable to account for inter-sample variability. GSEA was performed using the fgsea R package to compare pathways enriched between aAP LE GBCs and no-AP LE GBCs. The ranked gene list resulting from differential gene expression analysis was used as input. As a query, hallmark gene sets were obtained from the Molecular Signatures Database (MSigDB v2024.1). Cell–cell interaction analysis was performed using CellChat R package version 2.1.2^29^. The standard CellChat pipeline was used, with the cells split into aAP and no-AP, and the communication probabilities were calculated using triMean.

### DNA extraction

High molecular weight DNA from tumor tissue was extracted using Gentra Purigene Cell kit (Qiagen, 158767) following the manufacturer’s instructions. Briefly, 60 mg of tissue was lysed with 3 ml of Lysis Solution, followed by Proteinase K incubation overnight at 55 °C and RNase treatment for 5 min at 37°C. After 3 min incubation on ice, 100 µl Protein Precipitation Solution was added to the lysate to remove proteins, and then centrifuging for 1 min at 15,000 g. The supernatant was transferred into a clean 1.5 ml microcentrifuge tube with 300 µl isopropanol. DNA was visible as a white pellet after centrifugation. A final wash with 300 µl of 70% ethanol was done to remove remaining impurities. DNA was resuspended in 100 µl DNA Hydration Solution and incubated at 65°C for 1 h to dissolve the DNA. DNA concentration was measured by Qubit dsDNA BR Assay Kit (Thermo Fisher Scientific, Q32850), purity by Denovix spectrophotometer (DS-11Fx) and DNA integrity by TapeStation genomic DNA reagents and ScreenTape assay (Agilent 5067-5366 and 5067-5365, respectively).

### Shallow whole-genome sequencing (sWGS) library preparation and copy number estimation

Libraries were prepared using 100 ng of genomic DNA, following the NEBNext Ultra II FS DNA Library Prep Kit for Illumina (New England Biolabs, E7805S). Briefly, DNA was enzymatically fragmented for 19 min at 37 °C to get fragments averaging 450 bp. After size selection, end repair, dA-tailing and UDI Adaptor Ligation, final libraries were amplified by 5 PCR cycles and quantified by Qubit dsDNA High Sensitivity Assay Kit (Thermo Fisher Scientific, Q32851). Size distribution was assessed by Bioanalyzer High Sensitivity DNA Kit (Agilent, 5067-462). Libraries were combined at 4 nM concentration pool and sequenced paired-end on NextSeq 500 sequencer using NextSeq 500/550 Mid-Output v2.5 Kit (Illumina, 20024904), corresponding to 1X genomic coverage. Copy number variations (CNVs) were inferred from sWGS data using the Absolute Copy number Estimation (ACE) algorithm^30^.

### Paraffin Embedding and Hematoxylin and Eosin (H&E) Staining

See Supplementary Methods.

### Immunohistochemistry

Resliced FFPE slides were placed at 60 °C for 1 hour, followed by deparaffinization in Histolab-Clear (3 × 15 min), and sequential rehydration in absolute, 96%, and 70% ethanol. Slides were rinsed with distilled water and then 0.01 M PBS for 5 minutes. Heat-induced epitope retrieval (HIER) was conducted in a steamer preheated to 99.9 °C using Tris-EDTA buffer (pH 9) for 20 min. Slides were encircled with a PAP pen (Vector Laboratories), washed in PBS-T (PBS with Tween-20) for 5 min, and blocked with 0.2% skim milk in PBS-T for 30 min. Primary antibodies were diluted in PBS-T and applied individually after removing the blocking buffer. Incubation proceeded overnight at 4 °C in a humidified box. The following day, slides were washed in PBS-T (3 × 5 min) and incubated for 2Lhours at room temperature in the dark with species-specific secondary antibodies. After washing in PBS, nuclear counterstaining was performed by DAPI (1:10,000, 5 min). Slides were mounted with Fluoromount™ (Sigma-Aldrich), coverslipped, sealed with nail polish, and dried overnight. Prepared slides were stored in light-protected slide boxes at 4 °C before imaging.

### Antibodies and Morphology Reconstruction

Primary antibodies: rabbit anti-NeuN (Sigma-Aldrich, 1:1000) and mouse anti-Ki67 (Abcam, 1:1000). Secondary antibodies: donkey anti-rabbit Alexa Fluor 594 and donkey anti-mouse Alexa Fluor 488 (both 1:500, Abcam). Patched LE cells were reconstructed from stacks of 24–43 images using Neuromantic 1.7.5 software^31^.

### Statistical Analysis

Statistical analyses were performed in R (version 4.1.0) or GraphPad Prism 6 (GraphPad Software, United States). Data are presented as mean ± SEM unless otherwise indicated. Comparisons between two groups were conducted using unpaired two-tailed Student’s *T*-tests. For comparisons involving more than two groups, one-way ANOVA was used, followed by Tukey’s post hoc multiple comparisons test. When evaluating repeated or paired measures across three groups, a two-way ANOVA followed by Tukey’s multiple comparison test was applied. Correlations of electrophysiological parameters were assessed with linear regression and Pearson’s correlation. The classification of APs and aAPs was established by assessing the distribution of individual electrophysiological parameters through a normality test, followed by hierarchical cluster analysis. The identification of significant changes in gene expression was performed in R using MAST^32^. Levels of statistical significance are indicated as follows: * p<0.05, ** p<0.01, *** p<0.001, and **** p<0.0001.

## Results

We investigated tumor-infiltrated samples of neocortical brain tissue obtained from glioblastoma patients during fluorescence-guided neurosurgical resection (Figure 1A; Supplementary Figure 1). The experimental workflow involved the preparation of acute brain slices and AAV-mediated cell labeling in cultured slices (Figure 1B). To identify regions of tumor infiltration, H&E and immunohistochemistry staining were performed on adjacent slices (Figure 1C). Whole-cell patch-clamp recordings were performed to obtain electrophysiological and morphological readouts from cells located at the tumor LE or non-tumor-infiltrated cortex in both acute and cultured slices (Figures 1D-F). In a subset of recordings, we performed patch-seq by harvesting the cytoplasm and nuclei for single-nucleus RNA sequencing and analysis (Figures 1E-G). Other neighboring unpatched cells were also collected to increase yield for transcriptomic analysis (Figure 1G). The patched cells were loaded with intracellular dye to enable morphological reconstruction (Figure 1H).

**Figure.1.**
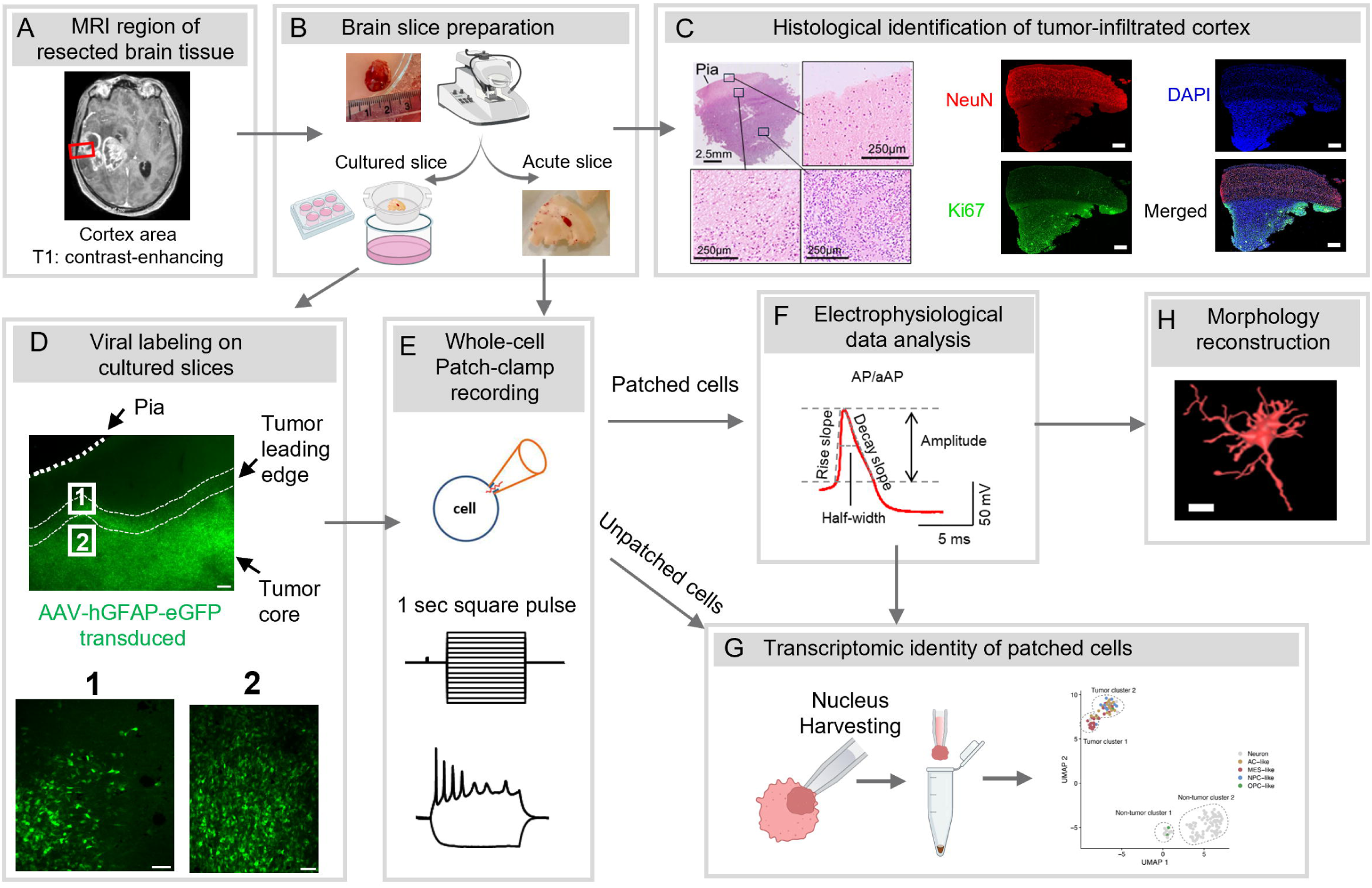
Workflow for utilizing patient-derived tumor-infiltrated neocortical tissue. **A**, The red box highlights the tumor-infiltrated neocortical with contrast-enhanced region on T1 MRI for neurosurgical resection. Tissue from this region was used for downstream *ex vivo* experimentation. **B**, Schematic overview of tissue processing. Surgically resected glioblastoma tumor-infiltrated cortical specimens were prepared for acute and organotypic slice cultures using a vibratome. Slices were either used as acute preparations for immediate patch-clamp recordings or maintained in culture for subsequent viral transduction. **C**, A representative image showing hematoxylin and eosin (H&E) staining of a tumor-infiltrated cortical brain slice from a glioblastoma patient. The disrupted cortical architecture and increased cellular density indicate tumor infiltration. Immunohistochemistry showing the target marker with NeuN (neuronal marker), DAPI (nuclei marker), and Ki67 (proliferation marker) in tumor-infiltrated human neocortical slices. Co-staining confirms the presence of infiltrating tumor cells within cortical tissue (scale bar: 500 µm). **D**, Cell-type-specific viral transduction using adeno-associated virus (AAV) vectors. Example demonstrating the use of the GFAP promoter to target astrocytes or glioblastoma cells (GFAP-eGFPD), allowing for fluorescence-based identification of labeled cells. **E**, Whole-cell patch-clamp recordings were performed on either acute slices or fluorescence-guided cultured slices to characterize the intrinsic membrane properties of identified cell types. **F**, Electrophysiological data were compiled for joint analysis using established parameters of intrinsic membrane properties. **H**, Seven patched cells were filled with Alexa Fluor 594 during recordings. Confocal z-stacks were acquired for 3D morphological reconstruction of the recorded cells, allowing detailed structural analysis. **G**, Following successful electrophysiological recordings, the nucleus of the patched cell was aspirated into the pipette and processed for single-nucleus RNA sequencing. To enhance transcriptomic coverage, nuclei from neighboring unpatched cells were also collected. The resulting transcriptomic data were analyzed to correlate intrinsic excitability with gene expression profiles.

### Neocortical LE cells display aberrant action potentials

To investigate electrophysiological and morphological properties, we performed whole-cell patch-clamp recordings targeting LE cells and non-LE neurons in the adjacent non-tumor-infiltrated neocortex. The LE was defined as the region infiltrated by tumor cells, located immediately superficial to the dense tumor core and adjacent to the structurally intact, preserved neocortex (Figure 2A). Pyramidal neurons located outside the tumor LE, referred to as non-LE neurons, were identified under bright-field microscopy at 40x magnification based on their characteristic triangular-shaped somata, measuring approximately 20–50 µm in diameter (Figure 2A, red inset)^33^. In contrast, LE cells formed dense clusters within regions of disrupted cortical lamination and displayed relatively round somata, typically measuring 10–20 µm^11^ (Figure 2B, orange inset). The non-LE neurons in acute slices displayed normal overshooting APs (Figure 2C). Intriguingly, we found that 59.6% (28 out of 47) of LE cells exhibited aberrant action potential (aAP) when depolarized by a positive current injection (Figure 2D). AP and aAP differed in the waveform features such as amplitude, half-width, maximum rise slope, and minimum decay slope (Supplementary Figures 2A, B), followed by a hierarchical cluster analysis (Supplementary Figure 2C), and separated by a 2D plot of amplitude and decay slope (Supplementary Figure 2D)^25^.

**Figure. 2.**
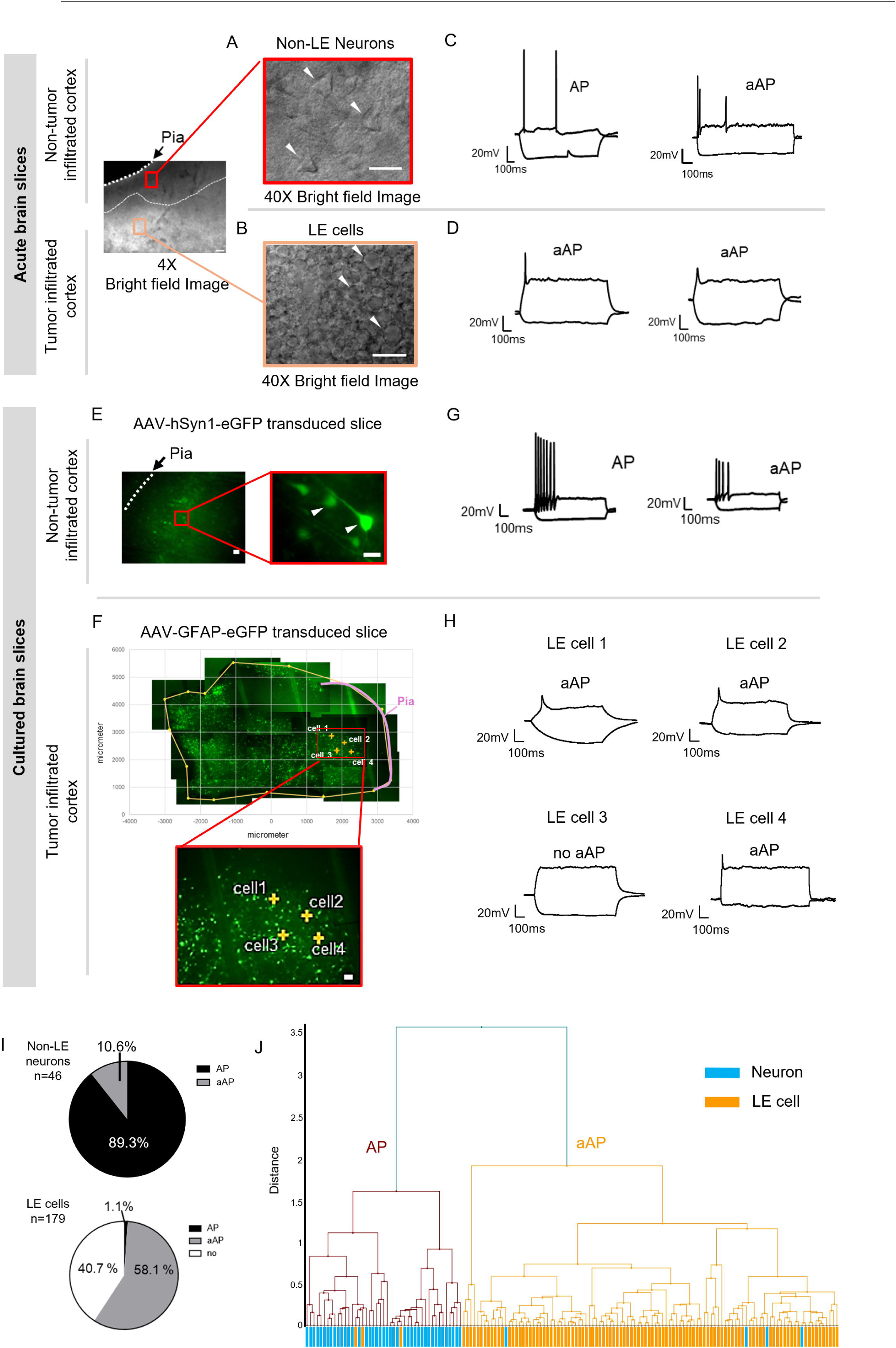
Neocortical tumor LE cells exhibit aAPs. Left upper panel: Bright-field image at 4x magnification showing a human tumor-infiltrated neocortical brain slice. The white dashed line delineates the boundary between preserved cortex (above) and tumor-infiltrated area (below). In the 4x magnification image, the scale bar corresponds to 100µm. In the 40× magnification image, the scale bar represents 20µm. **A**, High-magnification (40×) bright-field image of preserved neocortex (red inset) reveals pyramidal non-LE neurons with characteristic triangular somata (arrowheads). **B**, High-magnification (40×) bright-field image of the tumor LE cells (orange inset) shows dense clusters of small, round cells and disrupted laminar organization (arrowheads). **C**, Representative whole-cell patch-clamp recordings of neurons from acute slices showing one typical evoked action potential (AP) and one aberrant action potential (aAP). **D**, Representative whole-cell patch-clamp recordings from LE cells showing aAPs. **E**, Representative images from cultured brain slices show AAV-hSyn-eGFP labeling enabled visualization of non-LE neurons in the preserved neocortex. **F**, AAV-GFAP-eGFP labeling identified glial cells and putative glioblastoma cells (GBCs) at the tumor leading edge. **G**, Representative whole-cell patch-clamp recordings of AAV-hSyn-EGFP-labeled neurons in slice culture showed typical APs, with a minority exhibiting aAPs. **H**, Representative images show LE tumor cells 1–4, recorded from the neocortical tumor LE, exhibiting either aAPs or no excitability. **I**, Pie charts summarizing electrophysiological characteristics of all patched cells: 89.3% (42/47) of neocortical non-LE neurons exhibited APs, while 10.6% (5/47) showed aAPs. Among LE cells, 58.1% (104/179) exhibited aAPs, 40.7% (73/179) were non-excitable, and 1.1% (2/179) exhibited APs. **J**, Hierarchical clustering of electrophysiological parameters separated APs and aAPs across both non-LE neurons and LE cells. Note that a small fraction of non-LE neurons exhibited aAPs (4 cells), and 3 LE cells exhibited APs.

To target specific cell types of interest, we adopted an *ex vivo* slice culture paradigm combined with AAV-mediated enhancer-based fluorescent labeling^22–24^. We used AAV1-hSyn1-EGFP-WPRE-hGHp(A) to label neurons via the human synapsin-1 promoter (hSyn1), a conventional marker of neuronal identity^34^ (Figure 2E and red inset), and AAV5-hGFAP-hHEbl/E-EGFP-bGHp(A) to label GFAP-expressing glial cells, aiming for putative glioblastoma cells (GBCs) at LE (Figure 2F, red insets). hSyn1-eGFPL cells in structurally preserved cortex were identified as non-LE neurons based on their fluorescence, soma size, and evoked AP waveform (Figure 2E, G). In contrast, GFAP-eGFPL cells in the LE appeared as dense clusters of small, round cells within disrupted cortical structure, likely representing GBCs (Figure 2F).

The electrophysiological properties were examined by a standardized protocol testing active and passive membrane properties^35^. From all recorded cells in both acute and cultured brain slices (documented in Supplementary Figure 3), we observed that non-LE neurons were excitable upon positive current injection. Most non-LE neurons generated normal APs (89.3%, 42 out of 47 cells), with only five non-LE neurons exhibiting aAPs (10.7%, Figure 2I, J). Intriguingly, 58.1% (104 out of 179 cells) of total recorded LE cells exhibited active membrane properties upon depolarization (Figure 2H), whereas 40.7% (73 out of 179 cells) were non-excitable (Figure 2I). Only a small subset (1.1%, 2 out of 179) of LE cells displayed APs (Figure 2I, J). Morphological analysis of seven patched LE cells filled with Alexa594 and biocytin revealed structurally abnormal somatodendritic features, showing irregular processes (Supplementary Figure 4). The atypical morphology of LE cells is similar to that of previously reported neuron-like tumor cells observed in low-grade gliomas^36^. Collectively, these results demonstrate that the majority of LE cells exhibit neuronal-like excitability, characterized by the presence of aAPs, accompanied by morphological alterations.

### Distinct membrane properties and spontaneous excitatory input strength between neocortical non-LE neurons and LE cells

We compared the active membrane properties of non-LE neurons and LE cells in acute and cultured slices. Six AP/aAP features were measured, including: initiation (threshold, mV), amplitude (mV), half-width, maximum rise slope (mV/ms), maximum decay slope (mV/ms), and maximum firing rate (Figure 3A, B^37^). We found that LE cells exhibited a more depolarized threshold to initiate an aAP compared to non-LE neurons (Figure 3C, K). The aAP generated by LE cells had smaller amplitude, broader half-width, and slower rise and decay slopes compared to non-LE neurons (Figure 3D-G, L-O). Furthermore, LE cells fired fewer aAPs compared to non-LE neurons (Figure 3H, P). These features are reminiscent of immature neurons or early-stage stem cell-derived neurons^37, 38^. We also evaluated the consistency of active membrane properties over time in culture for both non-LE neurons and LE cells, and there was no significant trend of alteration of most electrophysiological features with increasing culture days (Supplementary Figure 5). In addition to membrane property measurements, in a subset of experiments, we recorded spontaneous excitatory postsynaptic current (sEPSC) from 40% of tested non-LE neurons (Supplementary Figure 6A) and 11% of tested LE cells (Supplementary Figure 6B). Non-LE neurons received more frequent sEPSC compared to LE cells, while LE cells showed faster sEPSC kinetics, with faster sEPSC onset and shorter half-width compared to non-LE neurons (Supplementary Figure 6C-H).

**Figure 3.**
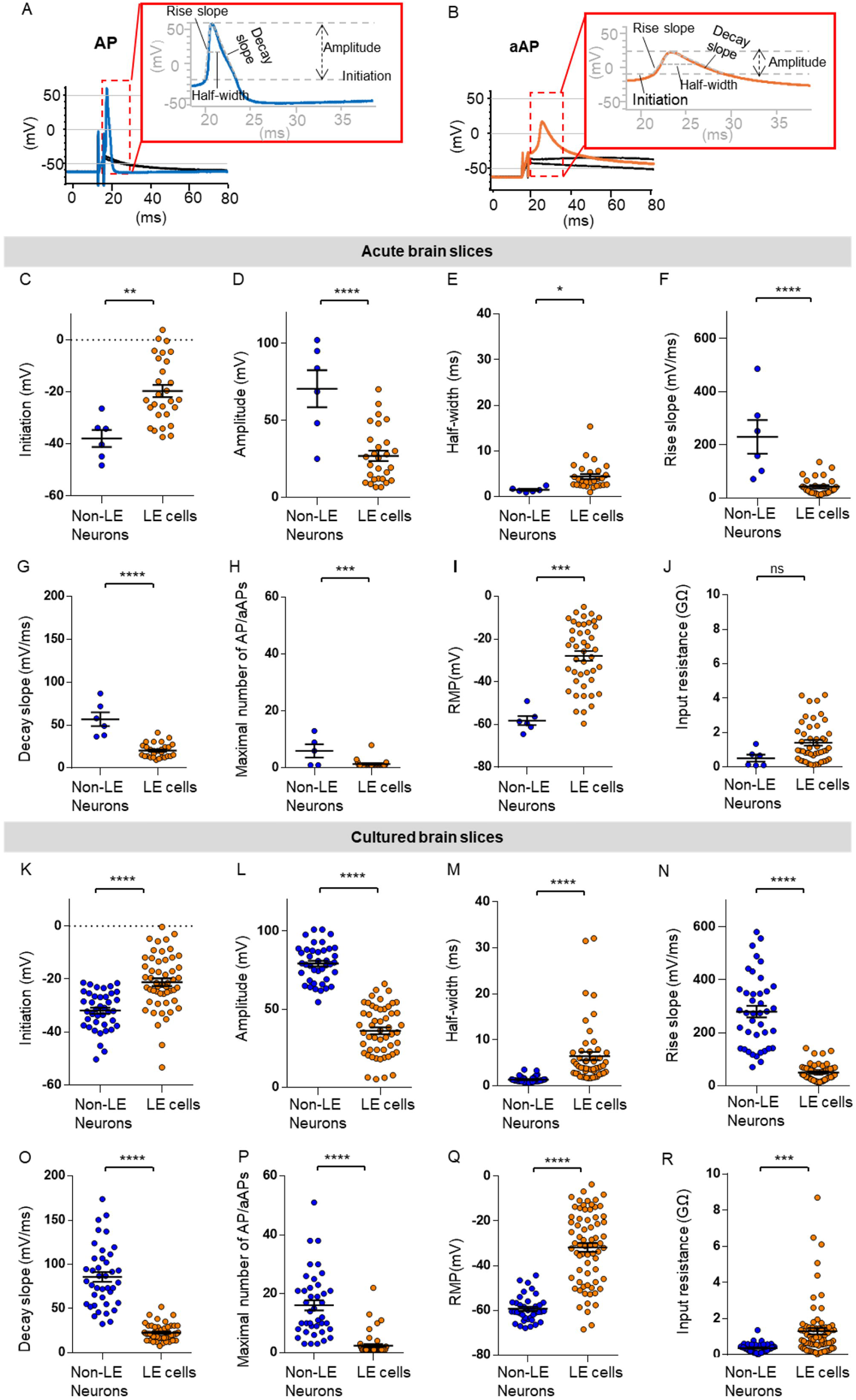
Electrophysiological characteristics of aAPs/APs in LE cells and neocortical non-LE neurons. **(A-B)** Representative traces showing an action potential (AP, **A**) and an aberrant action potential (aAP, **B**) evoked by a 3 ms depolarizing current injection. A schematic illustrates the key waveform parameters, including initiation, amplitude, half-width, rise slope, and decay slope, used to quantify AP and aAP properties. **(C–J)** Quantitative comparisons of APs/aAPs waveform parameters and passive membrane properties, including resting membrane potential (RMP) and input resistance, between non-LE neurons and LE cells recorded from acute brain slices. Boxplots show (**C**) initiation potential, (**D**) amplitude, (**E**) half-width, (**F**) rise slope, (**G**) decay slope, (**H**) maximal number of AP/aAP, (**I**) RMP, and (**J**) input resistance. In acute brain slices, non-LE neurons (blue, n = 6) exhibited significantly lower initiation potential (−37.95 ± 3.27 mV), higher amplitude (70.51 ± 12.05 mV), shorter half-width (1.51 ± 0.22 ms), faster rise slope (230.2 ± 63.03 mV/ms), faster decay slope (56.72 ± 8.07 mV/ms) and generated more APs/aAPs (6 ± 2.32) compared to LE cells (orange, n = 28) (initiation: −19.65 ± 2.35 mV, amplitude: 26.97 ± 3.38 mV, half-width: 4.40 ± 0.55 ms, rise slope: 42.63 ± 6.17 mV/ms, decay slope: 20.23 ± 1.55 mV/ms, maximal number of AP/aAP: 1.39 ± 0.26). Non-LE neurons displayed a more hyperpolarized RMP of −58.20 ± 2.14 mV and lower membrane input resistance of 0.52 ± 0.20 GΩ compared to LE cells (RMP: −27.84 ± 2.23 mV, input resistance: 1.41 ± 0.17 GΩ). **K–R**, Equivalent analyses from cultured brain slices. Boxplots show (**K**) initiation potential, (**L**) amplitude, (**M**) half-width, (**N**) rise slope, (**O**) decay slope, (**P**) maximal number of APs/aAPs, (**Q**) RMP, and (**R**) input resistance. In cultured slices, non-LE neurons (blue, n = 40) showed less depolarized initiation potential (−31.96 ± 1.15 mV), higher amplitude (79.14 ± 1.87 mV), shorter half-width (1.36 ± 0.11 ms), faster rise slope (279.7 ± 21.60 mV/ms), faster decay slope (85.65 ± 5.49 mV/ms) and generated more APs/aAPs (16.10 ± 1.73) than LE cells (orange, n = 52) (initiation: −21.19 ± 1.46 mV, amplitude: 36.10 ± 2.21 mV, half-width: 6.46 ± 1.05 ms, rise slope: 49.87 ± 4.30 mV/ms, decay slope: 22.58 ± 1.26 mV/ms, maximal number of AP/aAP: 2.39 ± 0.53). In cultured slices, non-LE neurons displayed a more hyperpolarized RMP of −59.34 ± 0.94 mV and lower membrane input resistance of 0.37 ± 0.04 GΩ than in LE cells (RMP: −31.88 ± 1.93 mV, input resistance: 1.30 ± 0.18 GΩ). Statistical significance: *p < 0.05, **p < 0.01, ***p < 0.005, ****p < 0.001.

In terms of passive membrane properties, the resting membrane potential (RMP) of LE cells was significantly depolarized compared to that of non-LE neurons (Figure 3I, Q). In addition, the input resistance (R_in_) of the LE cells was higher compared to non-LE neurons (Figure 3J, R). The RMP and R_in_ of all groups had no significant correlation with days *in vitro*, indicating stable passive membrane properties over the culture period (Supplementary Figure 5). These unique characteristics of LE cells, with relatively depolarized RMP and high R_in_, mimic those of hybrid cells recently identified in low-grade gliomas and set them apart from astrocytes, which typically have a much hyperpolarized RMP, lower membrane resistance, and are unable to generate aAP/APs^15^.

### aAPs generation in LE cells requires Na_V_ and K_V_ channels

To test if the ion channels required for LE cell aAP generation were similar to those of non-LE neurons, we first applied tetraethylammonium (TEA, 30 mM) to block voltage-gated potassium channels in general. This resulted in a notable decrease in the amplitude of the aAP compared to the baseline (Figure 4A-C). Additionally, we observed an increase in aAP half-width (Figure 4D) along with slower rise and decay slopes (Figure 4E, F). Subsequent bath application of tetrodotoxin (TTX, 1 μM), a voltage-gated sodium channel blocker, eliminated the aAP (Figure 4G). These findings elucidate the cellular components involved in LE cell aAPs and further confirm that the electrical excitability of neocortical LE cells bears similarity to that of immature neurons.

**Figure 4.**
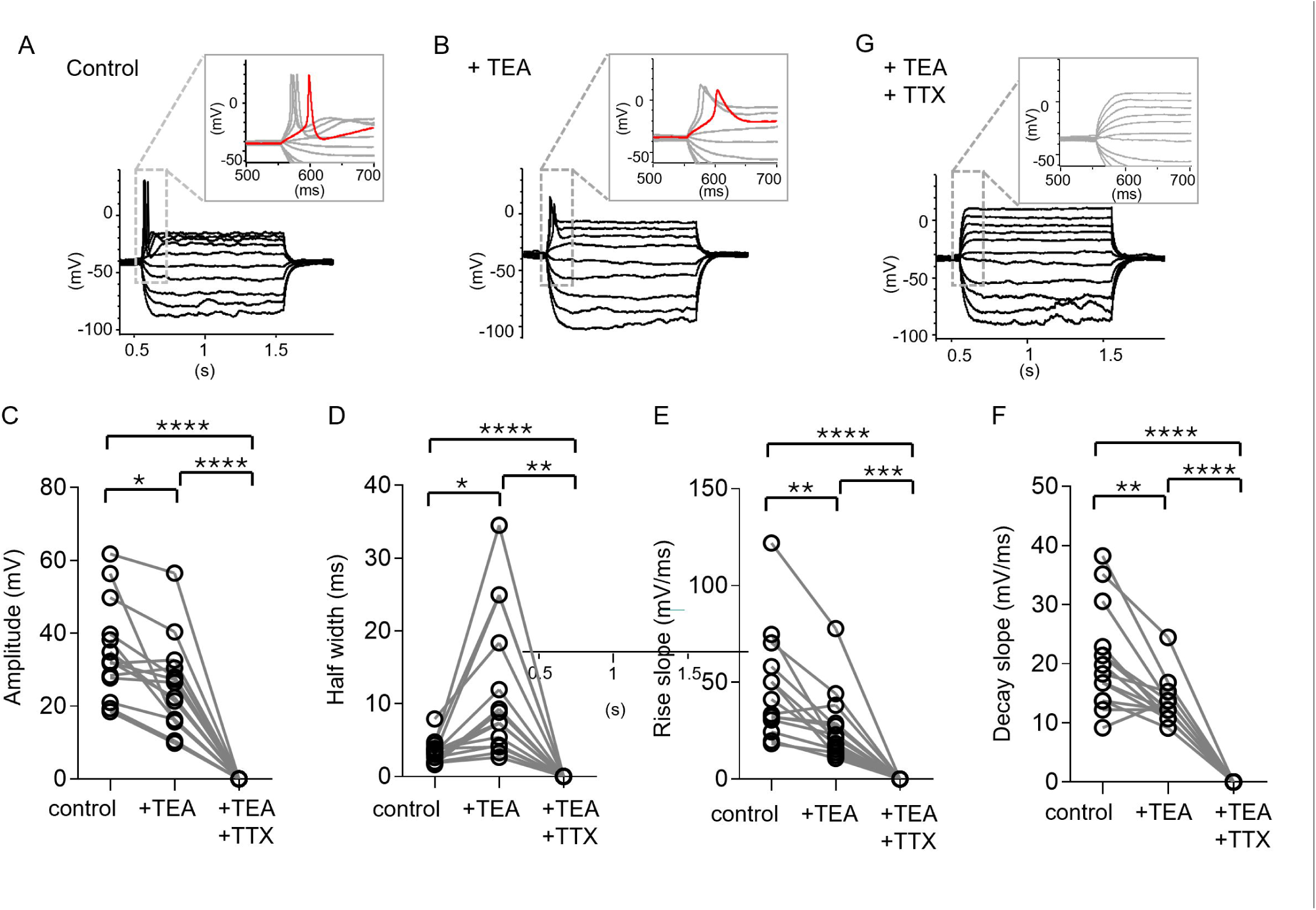
Contribution of Na_V_ and K_V_ Channels to aAP Generation. **A**, Illustrative aAPs induced by a 1-second square depolarizing pulse in a LE cell. **B**, Illustrations presenting changes in aAP morphology upon application of the K_V_ blocker TEA, highlighting alterations compared to the control condition. **C**, Comparison of amplitude parameter for 14 TME cells under three conditions: Control (35.13 ± 3.53 mV), TEA (25.32 ± 3.34 mV), and TEA+TTX (0 mV), exhibiting a reduction in aAP amplitude in the presence of TEA and a complete absence in the combined presence of TEA and TTX. TTX is a Na_V_ channel blocker. **D**, Comparison of half-width parameter under three conditions: Control (3.39 ± 0.43 ms), TEA (11.93 ± 2.64 ms), and TEA+TTX (0 ms), exhibiting a significantly wider half-width in the presence of TEA (* p < 0.05) and a complete absence in the combined presence of TEA and TTX. **E**, Comparison of rise slope parameter under three conditions: Control (46.97 ± 7.44 mV/ms), TEA (26.80 ± 4.70 mV/ms), and TEA+TTX (0 mV/ms), exhibiting a significant reduction in the presence of TEA and a complete absence in the combined presence of TEA and TTX. **F**, Comparison of decay slope parameter under three conditions: Control (20.77 ± 2.14 mV/ms), TEA (13.34 ± 0.94 mV/ms), and TEA+TTX (0 mV/ms), exhibiting a significant reduction in the presence of TEA and a complete absence in the combined presence of TEA and TTX. Comparative analysis among paired data from three groups was performed using two-way ANOVA followed by Tukey’s multiple comparison test. **G**, Absence of aAP induction upon depolarization after the combined application of both TEA and TTX. Statistical significance: *p < 0.05, **p < 0.01, ***p < 0.005, ****p < 0.001.

### Heterogeneous cellular states and distinct signaling in LE GBCs by patch-seq

To determine the cell types and molecular profiling of neocortical LE cells, we performed patch-seq experiments, which allow us to obtain both electrophysiological and transcriptomic data in LE cells under acute and slice culture configurations (Figures 1E-G; Figure 5A) [24]. To increase yields from each patient and improve transcriptomic mapping, we also harvested the LE cell nuclei for SMART-seq single-cell transcriptomic analysis only (Figure 5B; detailed cell information in Supplementary Table1). Collectively, a major subset (144 out of 154) of neocortical LE cells from 10 patients has qualified cDNA for further transcriptomic analysis, with 82 out of 144 cells having electrophysiological parameters measured (Figure 5B). We identified GBCs within those LE cells using a combination of label-transfer from a comparable dataset of glioblastoma tissue sampled at the tumor core and peripheral tissue^27^, and iterative inferring copy-number variance analysis, validated using shallow whole genome sequencing of the matched patient material^39^ (InferCNV; Figure 5C; Supplementary Figure 7; see Methods). The transcriptomic profiling of neocortical LE cells revealed four separated clusters, visualized in the 2D UMAP embedding (Figure 5D). As a result, two clusters of neocortical LE cells (53 out of 144 LE cells) were recognized as GBCs (Tumor clusters 1 and 2; Figure 5D), referred to as LE GBCs. In addition, the two clusters of non-tumor LE cells (non-tumor clusters 1 and 2) showed enrichment in neuronal transcriptomic profiling and hence were referred to as LE neurons (Darmanis dataset^27^; Figure 5E; Supplementary Figure 8A).

**Figure 5.**
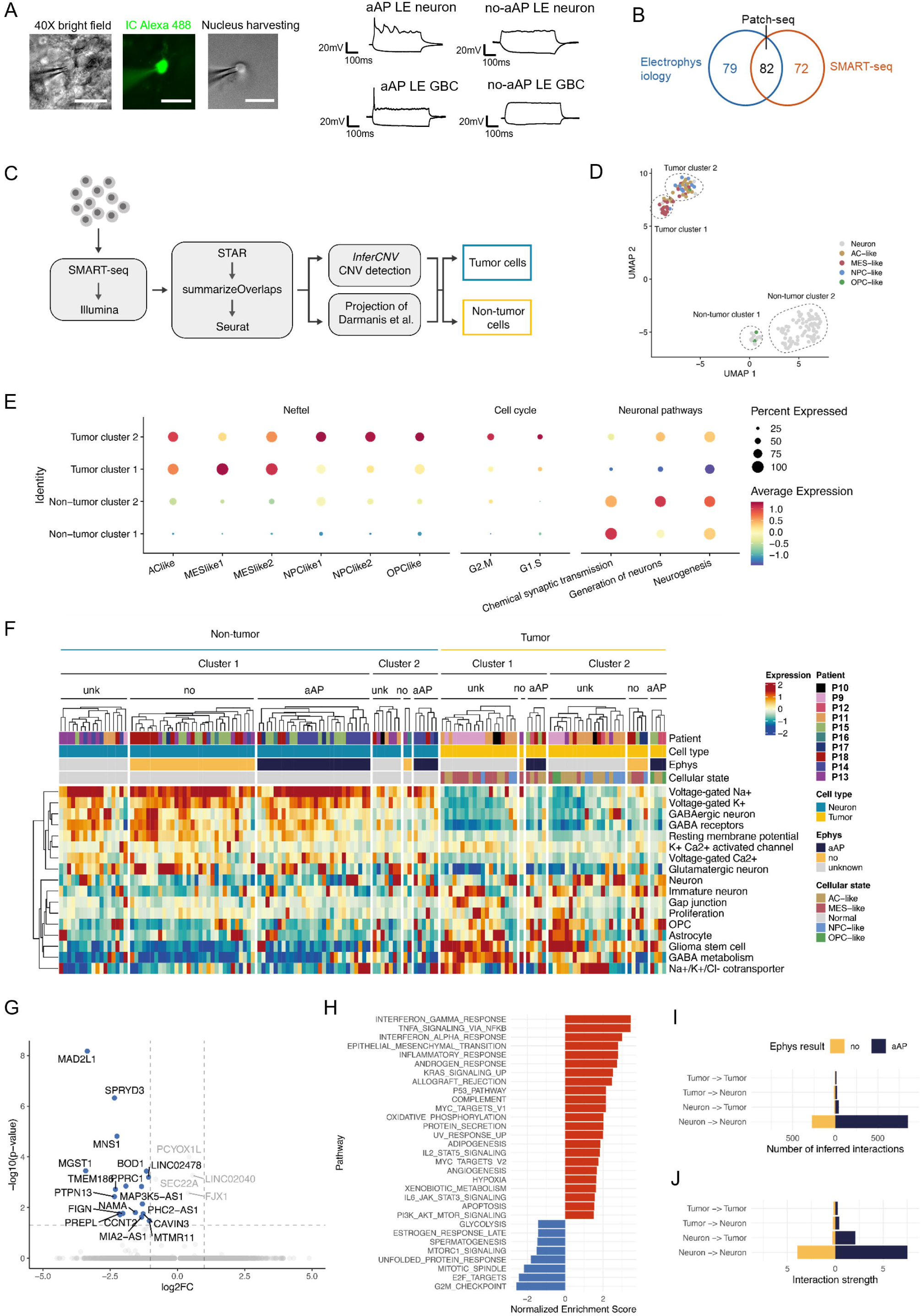
Characterization of aAP cells using Patch-seq and analysis workflow. A, Representative images of one cell that underwent the Patch-seq procedure (left), recorded with an intracellular solution containing Alexa 488 (middle). After electrophysiological recording, the nucleus was extracted (right) and collected for SMART-seq, followed by transcriptomic analysis. Representative electrophysiological traces from nuclei harvested from the neocortical tumor leading edge (LE), including aAP LE neuron, non-aAP LE neuron, aAP LE GBC, and non-aAP GBC. Scale bars: 20µm. **B,** Venn diagram illustrating the total number of cells included in electrophysiological and/or transcriptomic analysis. Out of 233 cells, 79 were analyzed only by electrophysiology, 82 had both electrophysiological and transcriptomic parameters measured, and 72 were collected only for transcriptomic analysis. **C,** Workflow of Patch-seq data generation and cell characterization. The single cells were processed using the SMART-seq protocol, followed by the identification of tumor cells and subsequent transcriptomic analysis. The resulting reads were aligned using STAR, counted using summarizeOverlaps, and imported into Seurat for downstream analysis. Cell type annotation was performed using a combination of InferCNV and cell type projection. **D**, UMAP visualization of cells, colored by their cellular state. The distinct tumor and non-tumor clusters are highlighted. **E**, A Dot plot showing the average expression of cellular state signatures, cell cycle, and neuronal pathways that were divided into four clusters. **F**, Heatmap of neuronal pathway expression across single cells. **G**, Volcano plot comparing gene expression between aAP LE GBCs and no-AP LE GBCs. **H**, Bar plot showing result of GSEA between the same groups as G. **I**-**J**, Bar plot showing cell-cell interaction results: (**I**) number of interactions, and (**J**) interaction strength.

As GBCs have heterogeneous cellular states, we next mapped the LE GBCs to the widely used cellular states defined by Neftel et al.^4^. The cellular composition was heterogeneous: one cluster predominantly consisted of MES-like cells (Tumor cluster 1, Figure 5D), while the other contained mainly AC-like and NPC-like cells (Tumor cluster 2, Figure 5D). Only five cells showed an OPC-like state (Figure 5A). In contrast, the LE neurons showed low expression levels of tumor cellular state markers (Figure 5E). In terms of cell cycle signaling, tumor cells, in general, and those in Tumor Cluster 1, showed enrichment of gene expression in the G1/S and G2/M cell cycle phases (Figure 5E). As expected from their cell type origin, the two transcriptomically identified LE neuron clusters exhibit enrichment of neuronal pathways compared to LE GBCs (Figure 5E).

15 out of 53 LE GBCs were electrophysiological assessed: 9 LE GBCs showed aAPs, while 6 LE GBCs did not generate aAP (no-aAP). 9 LE GBCs exhibiting aAPs were identified, including 2 MES-like, 5 AC-like, and 2 OPC-like cells. aAP GBCs can be sampled across patients and under both acute slice and slice culture configurations (Supplementary Figure 8B). On the other hand, 67 out of 91 LE neurons were electrophysiological assessed, with 34 LE neurons showing aAPs and 33 no-aAPs. The fraction of neurons and GBCs is similar under both acute and cultured slice configuration (Acute slice: 25 neurons, 5 GBCs; Cultured slice: 42 neurons, 10 GBCs). When comparing the electrophysiological properties of LE GBCs and LE neurons, we were surprised to find no significant difference in the electrophysiological properties between LE GBCs and LE neurons (Supplementary Figure 9). In contrast, transcriptomic profiling between LE GBCs and LE neurons revealed that LE neurons exhibited higher expression of Na_V_, K_V,_ and Ca_V_ ion channel signaling compared to LE GBCs (Figure 5E). On the other hand, LE GBCs exhibit stronger immature neuron and proliferation signaling, as well as other glial cell marker expressions (Figure 5F, Supplementary Figure 10A). Interestingly, unlike GBCs in the tumor core, the LE GBCs only showed weak expression of gap junction signaling-related genes (Figure 5F).

We next focused on the molecular properties of electrically distinct LE GBCs. We performed differential gene expression analysis followed by GSEA on the ranked gene lists between aAP and no-aAP LE GBCs. This analysis revealed significant enrichment in the mitotic pathway in aAP-negative compared to aAP-positive LE GBCs (Figure 5G, H). We also investigated the cellular states of electrically distinct LE GBCs. The cellular states of LE GBCs exhibited high heterogeneity in both acute and slice cultures, a phenomenon observed across patients, with slightly profound AC-like cell types in the LE (Supplementary Figures 8B-E)^4^. In terms of ion channel signaling, aAP LE GBCs showed a trend of higher expression of Na_V_ channel SCN3A compared to no-aAP LE GBCs, supporting their difference in aAP generation capability (Supplementary Figures 10F). Interestingly, in the LE, aAP-negative GBCs showed significantly higher expression of L-type voltage-gated calcium channel CACNA1C (Ca_V_1.2) compared to aAP-positive GBCs (Supplementary Figure 10F). Lastly, we applied CellChat to quantify the matched ligand-receptor expression between the LE GBCs and LE neurons, which showed a higher number of inferred interactions and higher interaction strength in neuron-to-neuron and neuron-to-tumor interactions within the aAP-positive compared to aAP-negative cells (Figures 5I, J).

In conclusion, patch-seq and transcriptomic analysis confirmed that LE GBCs can generate aAP with a similar electrical property as LE neurons, although several voltage-gated ion channel genes are more highly expressed in LE neurons. LE GBCs exhibited heterogeneous cellular states, and cells across different states displayed the capacity to generate aAPs, suggesting that excitability is a shared feature across different GBCs populations at the tumor leading edge. In addition, LE no-aAP GBCs were enriched for mitotic signaling, while LE aAP cells displayed stronger cell-to-cell interactions.

## Discussion

Glioblastoma is an incurable brain cancer characterized by significant cellular heterogeneity. Using human acute slices and slice cultures, combined with patch-seq, our study provides robust evidence of active electrophysiological features, heterogeneous cellular states with distinct proliferating features of GBCs at the LE. Moreover, LE neurons exhibited similar electrophysiological properties as GBCs, and aAP-generating cells demonstrated stronger ligand-receptor interactions in the LE.

We introduce an *in vitro* organotypic human brain slice culture model designed to facilitate long-term investigation of human tumor-infiltrated brain tissue using AAV-assisted labeling to target LE neurons and GBCs. Patch-seq confirmed that this model preserved key electrophysiological and molecular traits, with GBC-to-neuron sampling ratios similar to acute slices. By this approach, we found that about 58% of LE cells, including putative GBCs and neurons, both displayed aAPs with several similarities. Moreover, the aAP generation required voltage-gated Na^+^ and K^+^ channels. With patch-seq, we confirmed that transcriptomically and genomically defined LE GBCs were electrically active, displaying hyperexcitability and aAPs. In addition, LE GBCs exhibit enrichment of mitosis signaling compared to LE neurons; nevertheless, LE neurons display similar electrical properties to LE GBCs, despite their distinct molecular profiling compared to LE GBCs.

Both LE GBCs and LE neurons had depolarized resting potentials, high membrane resistance, and a tendency to fire single aAPs, unlike non-LE neurons. This may imply LE neurons exhibited an altered and aberrant electrophysiological phenotype, possibly due to tumor infiltration. The findings of hyperexcitability within the tumor-infiltrated region may likely be influenced by factors such as GBC-driven paracrine glutamate secretion^40,41^, reduced GABAergic inhibition^42^, and glioma-induced synaptic remodeling^43^. This aAP phenotype has also been documented under physiological conditions in neurons and certain glial populations^44–46^. Spikelets are observed in various neuron types, where they can arise from distal axonal action potentials, dendritic spikes, gap junctions, or ephaptic mechanisms^46^. In neurons, spikelets contribute to neuronal synchronization and modulation of synaptic plasticity^47, 48^. In glia cells, NG2-positive OPCs have been shown to exhibit aAP in the developing or injured central nervous system^44, 45^. Collectively, these findings suggest that aAP in LE GBCs may represent a specialized function potentially leading to active releasing mechanisms, invasion abilities, and local network integrations^40, 49, 50^, as proposed for neuron-like tumor cells in low-grade glioma^36^. Understanding the functional consequences of aAPs in LE GBCs may provide a new insight into the systemic tumor influence of nearby LE neurons.

The presence of aAP cells has not been observed in most cultured tumor cells or xenograft models (but see Sun et al. 2025^51^), and one plausible cause may be the different microenvironmental contexts. Cultured tumor cells and xenograft models^12–14^ may lack in vivo-like spatial, cellular, or vascular features as in acute slices, which could influence the electrophysiological phenotypes of GBCs. In contrast, studies reporting the excitability of GBCs have all utilized human glioblastoma slices^16–18, 36^, preserving the native cytoarchitecture, tumor-neuron interactions, and the tumor microenvironment. The study by Curry et al. (2024) identified hybrid cells (HCs) in gliomas that fire single brief action potentials and a transcriptomic profile combining GABAergic neuronal/ OPC-like features, including two HCs within *IDH1R132* wild-type patient samples^36^. Curry et al. reported the electrophysiological properties of the HCs closely resemble those of the aAP-generating cells observed in the current study. Our study specifically targeted the cortical LE cells from the transition zone between contrast-enhancing and non-enhancing regions on T1-weighted MRI with gadolinium contrast. Although the anatomical origin of the sample specimens in the Curry et al. (2024) study was not specified, our study demonstrates that such HCs, genomically defined as GBCs or neurons, are present at the cortical LE in all GBM patients examined. Moreover, aAP-generating LE GBCs were distributed across AC-like, MES-like, and OPC-like phenotypes, highlighting broader heterogeneity. This differs from the findings of Curry et al. (2024), in which all identified HCs in IDH-mutant gliomas were concurrently annotated as GABAergic- and OPC-like. Taken together, these results suggest that the capability of aAP generation is a general pathophysiological feature of glioma cells in both IDH-mutant and IDH-wildtype gliomas.

The aAP generation ability was linked to several cellular states of the LE GBCs. Such diversity may facilitate the creation of a communication framework with a maximized information encoding and processing capacity by assigning functional heterogeneous elements as much as possible, as has long been proposed in neuronal networks^52^. The diverse cellular and functional composition at the LE provides high flexibility in determining the fate and progression strategy of GBC cells based on the parameters in the dynamic tumor front line.

### Future Perspective

This study demonstrates the electrically active nature of GBCs in human glioblastoma, accompanied by the high molecular and electrical heterogeneity of the GBCs at the LE. Future studies using longitudinal activity monitoring of GBCs in human brain slice cultures from both the tumor core and LE, combined with specific viral tools for targeting GBC subtypes, will provide insights into understanding the detailed function of aAP in GBCs with different cellular states, and its link to tumor progression at distinct tumor compartments. In parallel, the mitosis-proliferation pathway is enriched in the LE no-aAP GBCs compared to the aAP GBCs, and further therapeutic attempts to knock down CCND2 on LE GBC progression need to be tested. Moreover, investigating the impacts of the tumor microenvironment and tumor-neuron interactions in relation to the alteration of physiological properties of neurons in the LE may reveal crucial mechanisms involved in GBM growth, invasion, and resilience, with the potential to develop targeted therapies.

### Limitations of Study

We acknowledge that our study is based on a limited number of patient samples, and that future studies will be needed to further elucidate the observed connectivity and heterogeneity in LE GBCs. In addition, there is a low level of voltage-gated ion channel gene expression detected from the patch-seq-harvested nuclei, especially in those identified as LE GBCs. Hence, although a trend is observed, we didn’t find a significant difference between aAP and no-aAP GBCs in the mRNA levels of voltage-gated ion channels that contribute to action potential generation. However, at the protein level, their electrical properties, in terms of the presence of voltage-gated Na^+^ and K^+^ channel functional properties, are distinct. The discrepancy between function and transcriptomic profiling may be due to 1. Cross-level comparison (gene vs. protein levels) and 2. methodological differences (electrophysiology measurement vs. transcriptomic molecular quantification), as well as technical limitations of sufficient mRNA material collection through the nucleus harvesting procedure for small-soma cells in patch-seq experiments^53^.

### Ethics

Human tissue procedures were approved by the Central Denmark Region Committee for Health Research Ethics (Journal Number: 1-10-72-82-17) and adhered to the World Medical Association Declaration of Helsinki principles.

## Funding

This study is supported by grants to AK (Danish Cancer Society R295-A16770 and R304-A17698-B5570; Aarhus University Research Fund AUFF-E-2023-9-41; Kræftfonden); to MC (Lundbeck-NIH grant number R325-2019-1490 funded by the Lundbeck Foundation; DFF-1 project 37741 funded by the Independent Research Fund Denmark) and to JDH (Krogh invest and Danish Cancer Society R295-A16770).

## Conflict of Interest

The authors declare no conflict of interest.

## Authorship

A.R.K. and M.C. conceived and conceptualized the project. T.T., W.-H.H., M.C., and A.R.K. designed the experiments. T.T. and W.-H.H. conducted and analyzed the electrophysiology experiments and performed the Patch-seq experiments. A.O. contributed to the electrophysiology experiments. J.D.H., F.G.R.G., and J.W. performed sWGS and Patch-seq transcriptomic data analysis, figure preparation, and manuscript writing. E.L.C. and J.T.E. contributed to immunofluorescence and morphology experiments. K.J.E. contributed to sample collection, metadata collection, and OncoPrint figure preparation. I.D.P., E.L.C., M.R., and K.P.F. contributed to human tissue transport, data entry, brain slice preparation, and organotypic slice culture maintenance. S.O.S.C., A.K.R.S., J.C.H.S., and A.R.K. contributed to patient inclusion and performed surgical procedures. K.M. provided data structure and management of patient data, and N.M. contributed to data entry and metadata. B.W.K. provided advice on immunostaining and histopathology experiments and contributed to manuscript editing. J.T.T. provided project advice and manuscript input. T.T., W.-H.H., and A.R.K. drafted the manuscript, and all authors contributed to critical editing. A.R.K. acquired funding and was responsible for project administration and management. A.R.K. and W.-H.H. supervised all aspects of the work.

## Data Availability

The data will be made available upon reasonable request.

## Acknowledgments

We thank K. Widzsiz for technical assistance and L.M. Fitting for logistical and technical assistance; Prof. Jens R. Nyengaard and T. W. Mikkelsen provided support for the histology experiment. H. Thygesen, A. Kobberø, and A.S. Møller-Andersen for administrative assistance; the OR staff at the Department of Neurosurgery, Aarhus University Hospital, for logistical assistance; E.M.R. Jensen for assisting patient enrolment. We thank the Bioimaging Core facility (RRID:SCR_023876) at the Department of Biomedicine at Aarhus University for providing access to instrumentation, training, and support. We thank the Danish Spatial Imaging Consortium (DanSIC), funded by the Novo Nordisk Foundation (grant NNF22OC0075296), for performing the immunohistochemistry slide scanning. We thank the Danish Neuro Single Cell (NeuSiC) platform at the Biotech Research and Innovation Centre (BRIC), University of Copenhagen, for library generation and sequencing the Patch-seq samples.

## Notes

### Competing Interest Statement

The authors have declared no competing interest.

### Summary of Updates

In this revised version, we have substantially updated the manuscript. We performed additional Patch-seq experiments with subsequent transcriptomic analyses to confirm our key observation that glioblastoma cells (GBCs) at the tumor leading edge (LE) are hyperexcitable and can generate aberrant action potentials (aAPs) under both acute and organotypic cultured human brain slices. These experiments directly address the major concern regarding the identity of aAP-generating cells. Using Patch-seq, we verified that molecularly defined GBCs at the LE, validated by shallow whole genome sequencing of matched patient material, generate aAPs in distinct patients. LE GBCs exhibit heterogeneous cellular states including astrocytic-like, mesenchymal-like and neural progenitor-like states, with a minority of oligodendrocyte precursor-like cells. aAP-generating GBCs showed lower expression of mitotic signaling compared to non-aAP GBCs, and CCND2 emerged as a transcriptomic marker distinguishing neurons from GBCs. We also analyzed neurons at the LE. Transcriptomically defined neurons at the LE generate pathological aAPs or fail to generate aAPs, with electrical properties indistinguishable from those of GBCs. Cells with aAPs at the LE show stronger cell-cell interaction signaling. Finally, we expanded the scope of the manuscript by adding measurements showing that GBCs at the LE receive excitatory postsynaptic currents with slightly faster kinetics than LE neurons. We also provide detailed transcriptomic profiling and analysis of key pathways such as mitosis signaling and cell-cell interaction strength for both GBCs and neurons.

