## Supplemental figures and supplementary method for "Glioblastoma cells imitate neuronal excitability in humans"

### Supplementary Methods

#### Paraffin Embedding and Hematoxylin and Eosin (H&E) Staining

Adjacent acute tumor-infiltrated neocortical brain slices were fixed in 4% paraformaldehyde (PFA), processed into paraffin blocks, sectioned at 3 µm, and mounted on adhesion slides. Sections were deparaffinized, rehydrated, stained with hematoxylin and eosin, dehydrated through graded ethanol, cleared in xylene, and coverslipped. After drying overnight, slides were imaged on a Hamamatsu NanoZoomer (C9600-12) at 40x magnification.

H&E staining was performed on paraffin-embedded tissue sections. Slides were deparaffinized and rehydrated following with two 20× dips in distilled water. Sections were stained with hematoxylin (vendor) for 10 minutes, washed under cold running tap water for 10 minutes, counterstained with eosin (vendor) for 2 minutes, and dehydrated through ethanol (20× dips in 70%, followed by 2 × 5 min in 96%, and 3 × 5 min in 99% EtOH). Clearing was achieved by 2 × 20 dips and 1 × 5 min incubation in xylene. Slides were then mounted with mounting medium and sealed with coverslips (Hounisen). After drying overnight, slides were imaged on a Hamamatsu NanoZoomer (model C9600-12) at 40x magnification.

### Supplementary Figures

**Supplementary Figure. 1**

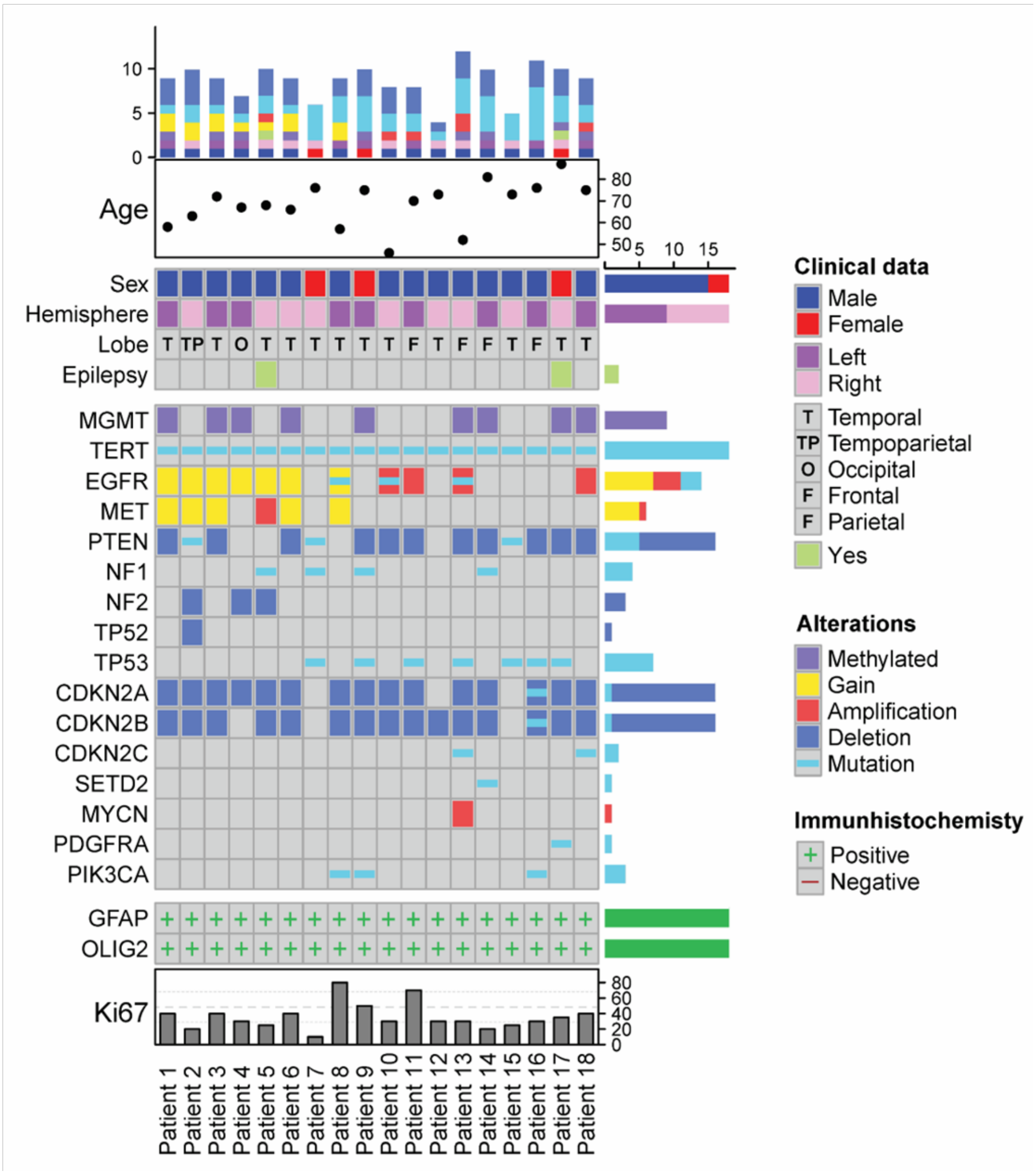


**Supplementary Figure 1** An OncoPrint plot showing a comprehensive visual representation of both the clinical profiles and genomic landscapes of eight glioblastoma patients. At the top of the print, age, sex, tumor location, and epilepsy are shown. In the middle part, an overview of genomic alterations (legend) in particular genes (rows) affecting individual samples (columns) can be seen. Different colors and shapes distinguish the type of alterations. The genes (rows) are sorted based on the frequency of the gene-level alterations in the cohort, as noted on the left of the figure. At the bottom of the print, immunohistochemistry analysis from the pathology reports is recapped. The percentage of Ki67-positive cells is illustrated by a bar chart. The bars at the top of the figure summarize the clinical characteristics and molecular landscape identified in each patient.

**Supplementary Figure. 2**

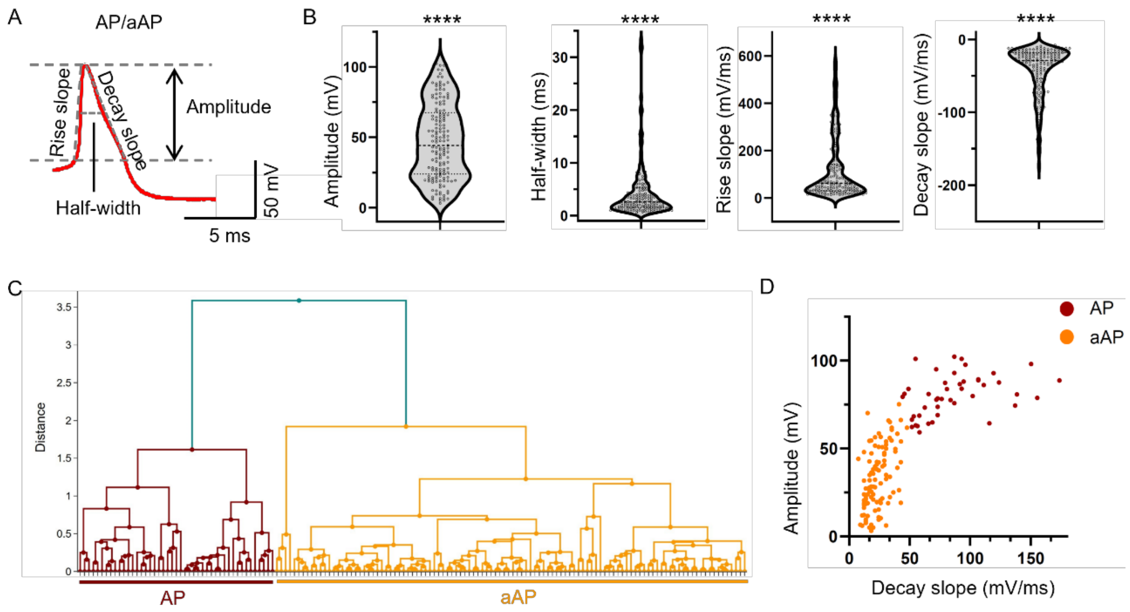


**Supplementary Figure. 2. Categorization of AP and aAP. A**, A schematic illustrates an AP/aAP evoked by a brief 3ms-depolarization pulse. The AP/aAP is characterized using four distinct parameters: amplitude, half-width, and rise/decay slopes. **B**, Violin plot of amplitude, half-width, rise slope, and decay slope from all recorded excitable cells. The statistical significance from the D'Agostino–Pearson omnibus normality test demonstrates that the parameters presented are not unimodally distributed. Asterisks indicate significant deviation from a normal distribution (****$p < 0.0001$). **C**, Hierarchical cluster analysis of all recorded excitable cells with electrophysiological parameters shown in B as the parameters for classification. The x-axis of the dendrogram represents the individual cell counts, and the y-axis represents the squared Euclidean distance between cells and clusters. **D**, Scatterplot of amplitude versus decay slope. Cells classified as AP and aAP phenotypes in the dendrogram are red and yellow, respectively.

**Supplementary Figure. 3**

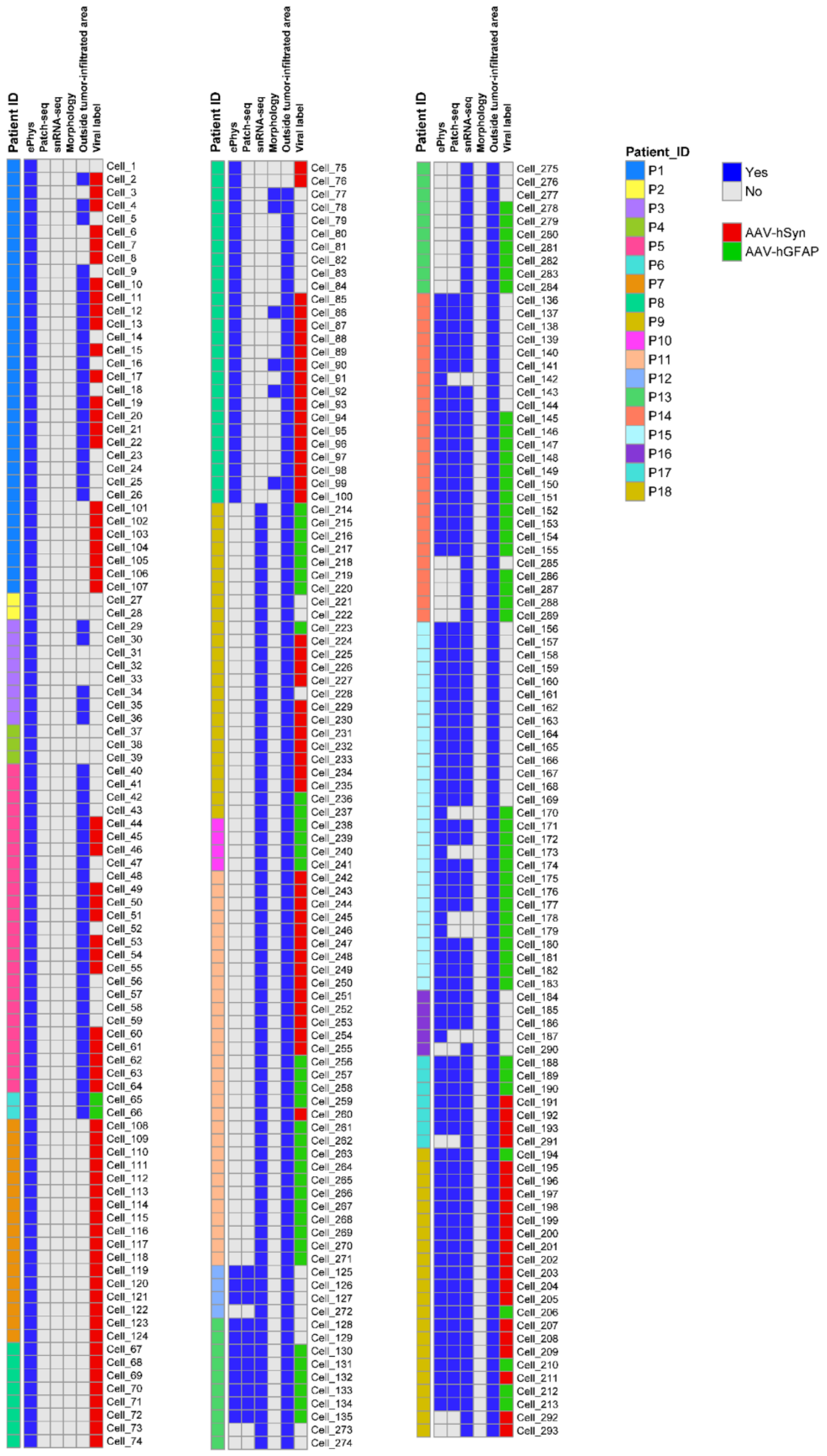

**Supplementary Figure. 3 Metadata overview of all recorded cells from glioblastoma patients.** The heatmap summarizes metadata for all 293 individual cells collected from 18 glioblastoma patients in this manuscript. Each row represents a single cell, annotated by cell ID (left column), and each column indicates experimental metadata, including tissue origin (outside tumor-infiltrated area or within the tumor leading edge), morphology reconstruction (Yes/No), electrophysiological recording (Yes/No), Patch-seq analysis (Yes/No), single-nucleus RNA sequencing (snRNA-seq), and viral labeling (AAV-hSyn-eGFP, marked in red; AAV-hGFAP-eGFP, marked in green). Patient identity is shown as color-coded bars on the far right. The metadata shows which experiments, electrophysiology, transcriptomics, and/or viral labeling, were successfully performed on each individual cell. This integrated overview underscores the heterogeneity of cell sampling and the multimodal nature of single-cell analysis across the patient cohort.

**Supplementary figure 4**

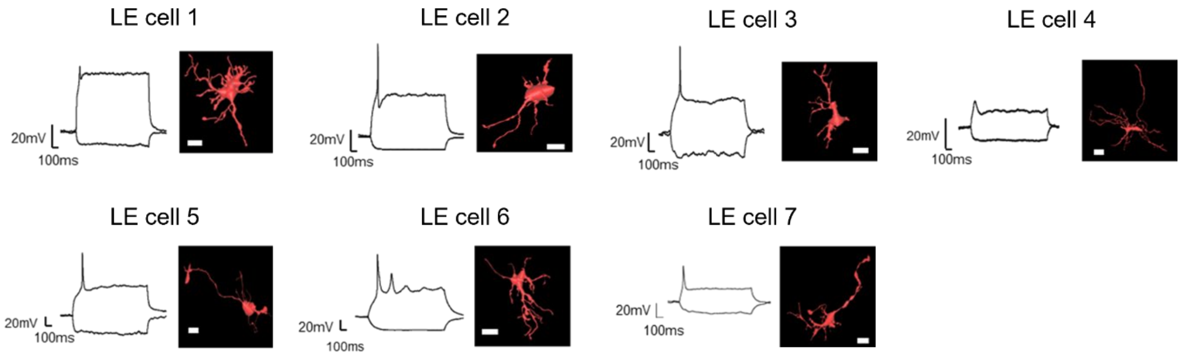


**Supplementary Figure 4. 3D morphology reconstruction of the patched LE cells.** Each reconstruction illustrates the somatodendritic architecture of individual cells recorded from the LE (7 cells in total) in human glioblastoma-infiltrated neocortical slices. Morphologies reveal variability in soma size, dendritic arborization, and process complexity across cells, highlighting the structural heterogeneity of LE cell populations. Scale bar: 20 μm.

**Supplementary Figure 5:**

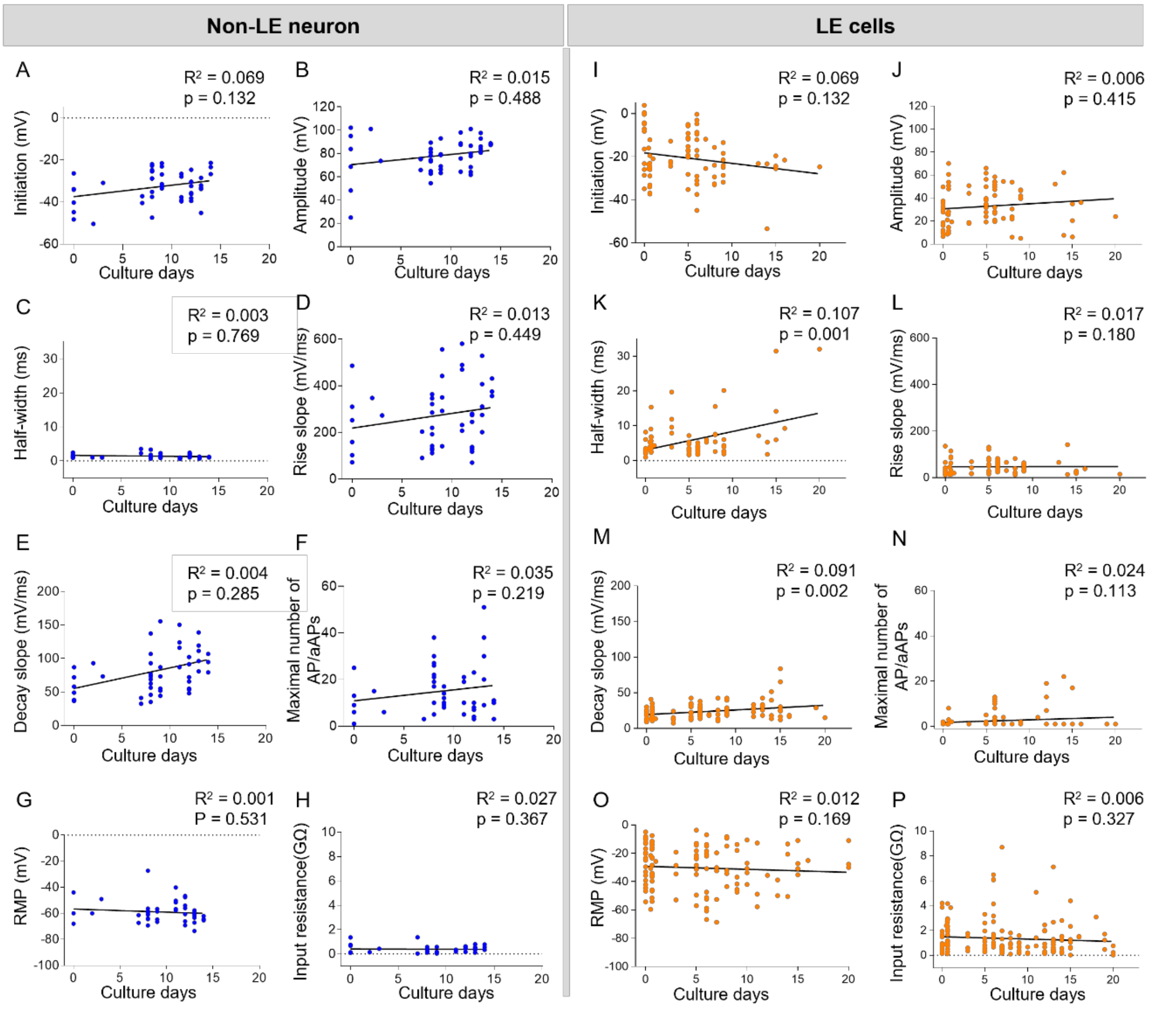


**Supplementary Figure 5: Dynamic analysis of electrophysiological parameters of AP/aAP over culture time in non-LE neurons and LE cells over the culture period.** The figure displays linear regression analyses of electrophysiological waveform parameters and passive membrane properties over time in culture for neocortical non-LE neurons (left, blue) and LE cells (right, orange). Parameters include initiation, amplitude, half-width, rise slope, decay slope, maximal number of AP/aAPs, resting membrane potential (RMP), and input resistance. Pearson's correlation and linear regression were used to assess trends over time. **A–H**, Non-LE neurons showed no significant correlation between culture time and any measured parameters (all $R^2 < 0.069$, $p > 0.05$). **I, J, L, M, N, O, P**, LE cells exhibited no consistent significant correlation between culture duration and electrophysiological or passive properties (all $R^2 < 0.065$, $p > 0.05$). **K** and **M**, for LE cells, there are weak correlations in threshold, amplitude, and half-width ($R^2 < 0.107$ and $p < 0.05$) over the culture period. These results suggest that both non-LE neurons and LE cells maintain relatively stable electrophysiological and intrinsic membrane properties during the *ex vivo* culture period.

**Supplementary Figure. 6:**

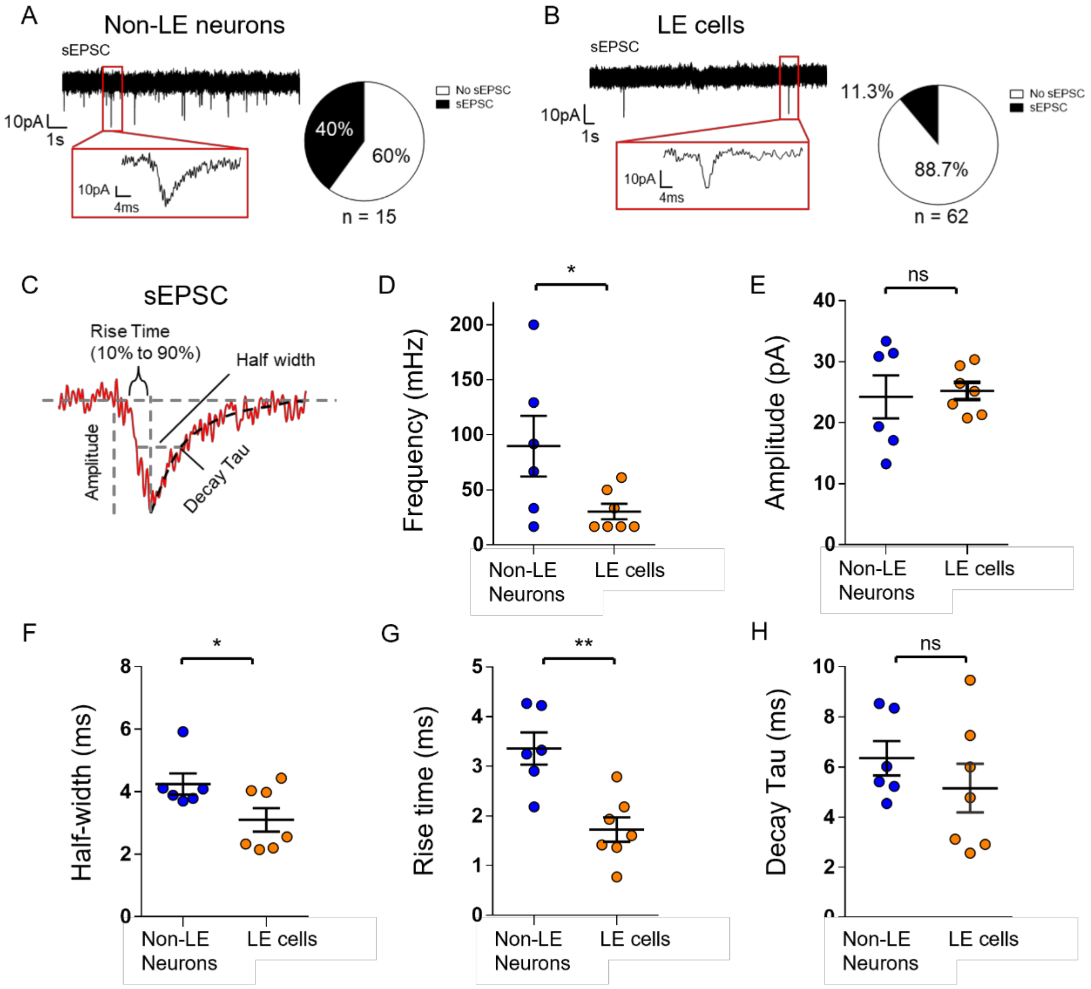

**Supplementary Figure 6. Comparison of excitatory synaptic input between neocortical non-LE neurons and LE cells.** (**A, B**) Representative sEPSC recordings from non-LE Neurons (**A**) and LE cells **(B)**. The insets showed a single EPSC event from the traces. The pie charts showed that **A,** a total of 40% of non-LE neurons had sEPSC detected (6/15 tested neurons: 2/4 neuron from acute and 2/11 neurons from slice cultures). **B**, Among LE cells, 11.3% had sEPSC detected (7/62 tested cells: 1/11 cells from acute and 6/51 cells from slice cultures). (**C**) Schematic illustration of the measured sEPSC properties, including amplitude, half-width, rise time (10%–90%), and decay tau. (**D–H**) Quantitative comparisons between neocortical non-LE neurons (blue) and LE cells (orange) of sEPSC properties. Non-LE neurons (n = 6) exhibited significantly (**D**) higher sEPSC frequency (89.58 ± 27.55 mHz), (**F**) slower half-width (4.25 ± 0.34 ms), (**G**) and longer rise time (3.36 ± 0.33 ms) compared to LE cells (n = 7) (sEPSC frequency: 30.16 ± 7.05 mHz, half-width: 3.10 ± 0.38 ms, rise time: 1.72 ± 0.25 ms). No significant differences were observed in (**E**) sEPSC amplitude (non-LE neurons: 24.22 ± 3.53 pA; LE cells: 25.20 ± 1.42 pA) and (**H**) decay tau (non-LE neurons: 6.35 ± 0.69 ms; LE cells: 5.15 ± 0.97 ms). Statistical significance: *$p < 0.05$, **$p < 0.01$.

**Supplementary Figure 7:**

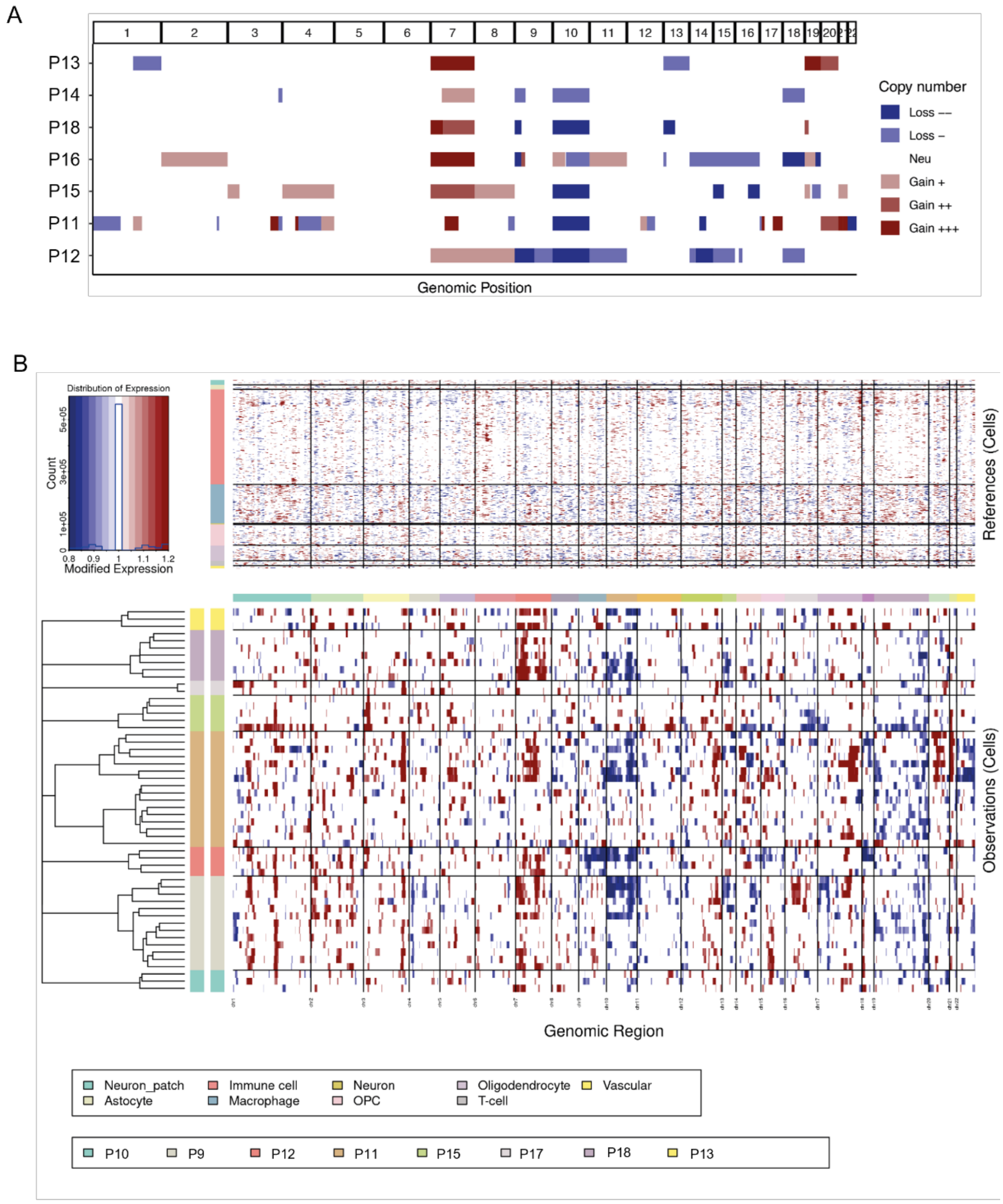


**Supplementary Figure 7.** Copy-number characterization of LE GBCs. A, Genome-wide copy-number profiles of each patient, derived from shallow WGS using ACE [38]. Colors of the segments denote the copy number status. B, Single-cell inferred CNV of LE GBCs across patients, generated using InferCNV. The upper panel illustrates the normal cells used as reference, including neuron cells from our patch-seq LE dataset and additional normal cell types from the Darmanis et al. dataset [27]. The lower panel displays the LE GBCs, with the color indicating the inferred copy number level within a cell.

**Supplementary Figure 8:**

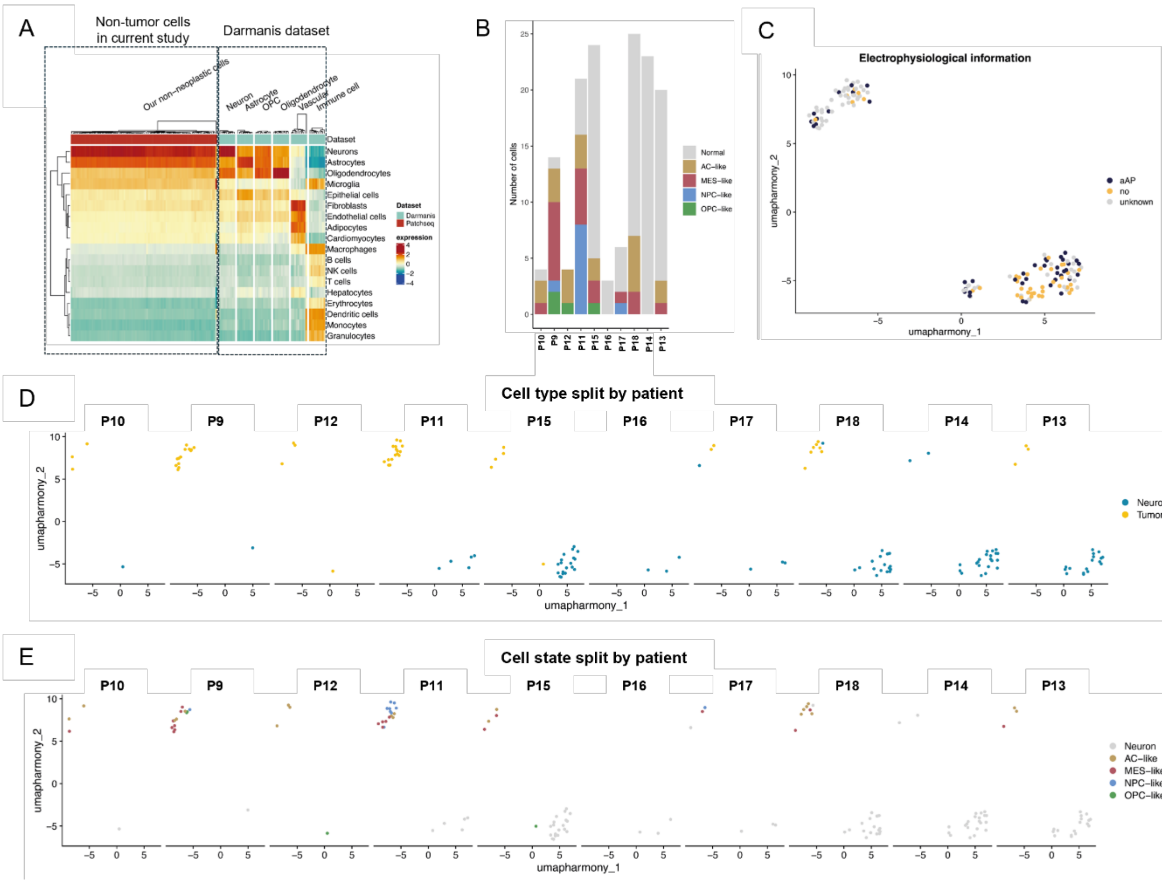


**Supplementary Figure 8. Transcriptomic and electrophysiological characterization of neocortical LE cells. A**, Heatmap showing label transfer of LE cells to reference cell types from the Darmanis dataset, confirming the presence of non-neoplastic neuronal identities, except one microglia cell identified. **B**, A barplot of inferred cellular states across patients, including AC-like, MES-like, NPC-like, OPC-like, and non-tumor neurons. GBCs with different cell states can be sampled within the same patient, and this phenomenon is consistent across patients. **C,** UMAP embedding of LE cells with electrophysiological annotation, showing the distribution of cells that generate aAP, cells without aAP (no-aAP), and nucleus-harvested-only cells with unknown electrophysiological profiles. **D**, UMAP plots showing cell type distribution (tumor vs. neuron) separated by patient. **E**, UMAP plots showing inferred cellular states of LE cells separated by patient, highlighting heterogeneous tumor cell states (AC-like, MES-like, NPC-like, OPC-like) compared with neurons.

**Supplementary Figure 9:**

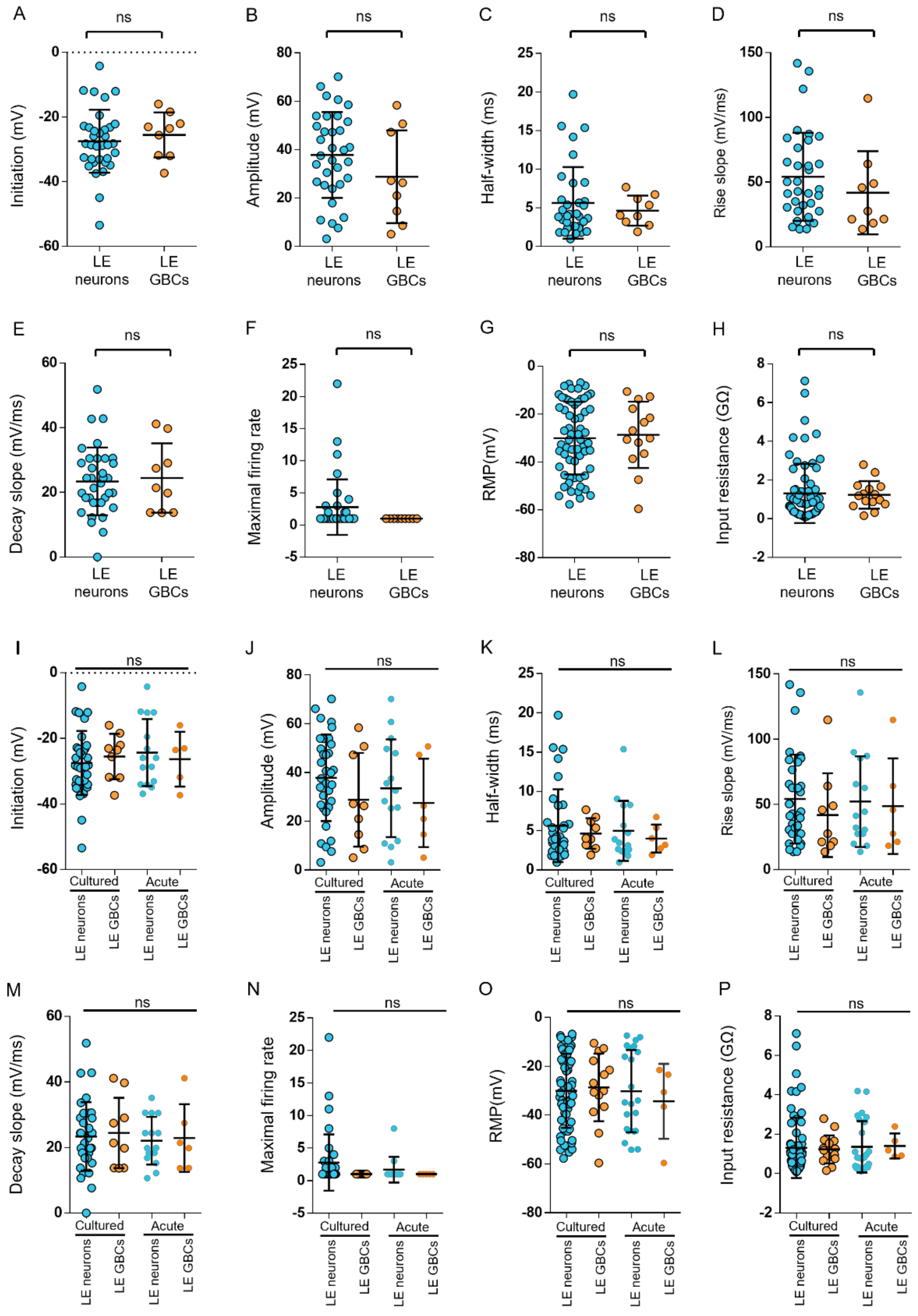


**Supplementary Figure 9. Electrophysiological comparisons between transcriptomically identified LE neurons and LE GBCs. A–H,** Quantitative analysis of electrophysiological properties of Patch-seq recorded cells, classified as LE neurons or LE GBCs based on single-nucleus transcriptomic profiles. Electrophysiological parameters include (**A**) initiation potential, (**B**) amplitude, (**C**) half-width, (**D**) rise slope, (**E**) decay slope, (**F**) maximal number of APs/aAPs, (**G**)

RMP, and (**H**) input resistance. No significant differences were observed between LE neurons or LE GBCs in all the measured parameters. **I–P,** Analysis of LE neurons or LE GBCs recorded from both acute and cultured brain slices. Electrophysiological parameters include (**I**) initiation potential, (**J**) amplitude, (**K**) half-width, (**L**) rise slope, (**M**) decay slope, (**N**) maximal number of APs/aAPs, (**O**) RMP, and (**P)** input resistance. There is no significant difference between LE neurons or LE GBCs in both acute and cultured conditions.

**Supplementary Figure 10:**

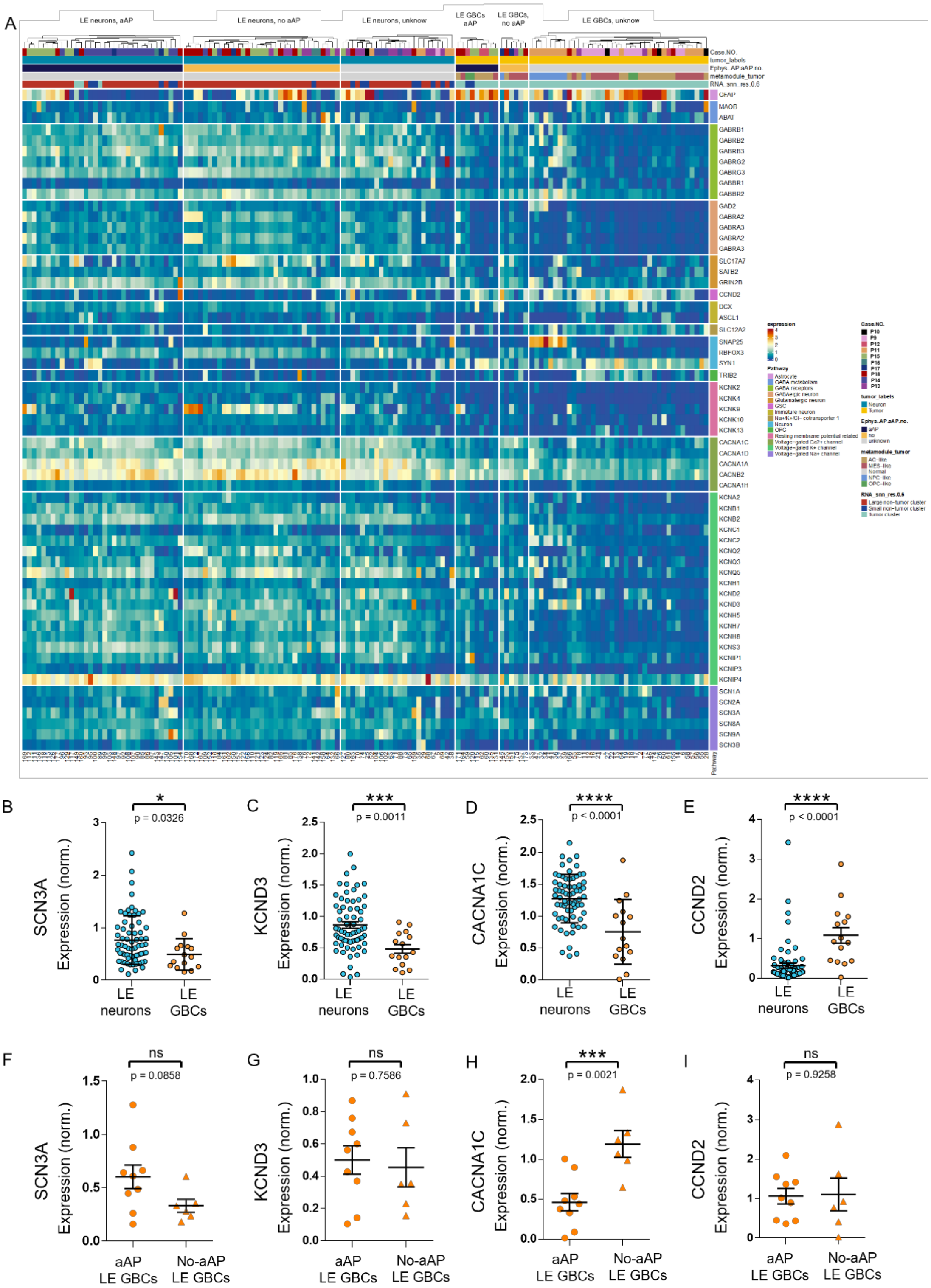


**Supplementary Figure 10. Transcriptomic and electrophysiological correlation of ion channel gene expression in LE neurons and LE GBCs. A,** Heatmap showing the expression of selected ion channel genes in Patch-seq samples. Hierarchical clustering separates LE neurons and LE GBCs into distinct groups, including neurons with aAP (LE Neurons-aAP), LE neurons without excitability (LE Neuron-no-aAP), LE GBCs with aAPs (LE GBCs-aAP), and non-excitable tumor cells (LE GBCs-no-aAP). Gene families include GABA receptors, glutamate receptors, potassium channels, calcium channels, and sodium channels. Color scale indicates normalized expression levels. **B–E,** Comparison of the expression of selected genes (**B**) SCN3A, (**C**) KCND3, (**D**) CACNA1C, and (**E**) CCND2 between LE neurons (blue circles) and LE GBCs (orange symbols) with available electrophysiological recordings. **F–I,** Comparison of the expression of selected genes (**F**) SCN3A, (**G**) KCND3, (**H**) CACNA1C, and (**I**) CCND2 between aAP- (orange circles) and no-aAP LE GBCs (orange triangle). Statistical significance: *$p < 0.05$, **$p < 0.01$, ***$p < 0.005$, ****$p < 0.001$.
